# CENP-A and CENP-B collaborate to create an open centromeric chromatin state

**DOI:** 10.1101/2022.07.08.499316

**Authors:** Harsh Nagpal, Beat Fierz

## Abstract

Centromeres, the sites within chromosomes responsible for accurate genome repartitioning, are epigenetically defined via replacement of canonical histone H3 by the histone variant CENP-A forming specific nucleosomes with increased DNA flexibility. In human cells, CENP-A nucleosomes and thus centromeres localize to genomic regions containing extended tandem repeats of alpha-satellite DNA. There, the constitutive centromere associated network (CCAN) and the kinetochore assemble, connecting the centromere to spindle microtubules during cell division. CENP-A provides a major recruitment point for many CCAN member proteins. One factor, CENP-B, binds to a specific DNA sequence contained in about half of alpha-satellite repeats. CENP-B is a dimer and is involved in maintaining centromere stability and, together with CENP-A, shapes the basic layer of the centromeric chromatin state. While recent studies have revealed the structure of large parts of the CCAN complexes, the nanoscale organization of centromeric chromatin is not well understood.

Here, we use single-molecule fluorescence resonance energy transfer (FRET) and colocalization imaging as well as dynamic experiments in cells to show that CENP-A incorporation establishes a far more dynamic and open chromatin state compared to canonical H3. We investigate whether CENP-A marks a landing spot for CENP-B, and find that on the single nucleosome level, CENP-B does not prefer H3 over CENP-A nucleosomes. However, in a chromatin fiber context, CENP-B binding is suppressed by higher-order chromatin structure. The increased dynamics of CENP-A chromatin create an opening, allowing CENP-B access and binding. In turn, bound CENP-B further opens the chromatin fiber structure, potentially via bending the bound DNA. Finally, transient knockdown of CENP-A expression in cells increases CENP-B mobility in cells. Together, our studies show that the two centromere-specific proteins collaborate to reshape chromatin structure, enabling the binding of centromeric factors and establishing a centromeric chromatin state.

## Introduction

Chromatin, the nucleoprotein complex composed of histone proteins and genomic DNA, is an essential regulator of processes such as gene expression and DNA replication, as it controls both local and gene-scale genome organization and accessibility^1–3^. The chromatin state is defined by local deposition of histone protein variants^4^, as well as histone post-translational modifications, and by the local accumulation of a plethora of different chromatin factors^5^.

CENP-A, the centromeric histone H3 variant acts as the epigenetic marker for centromere formation and subsequent DNA segregation^6–8^. CENP-A-containing nucleosomes are deposited on AT-rich DNA^9,10^ within centromeric chromatin (centro-chromatin), which consists of tandem repeats of alpha-satellite DNA in human cells^11^. Within these centromeric regions, CENP-A acts to recruit the kinetochore^12–14^, a large protein complex required to bridge the chromosomes with the spindle microtubules. CENP-A exhibits a number of structural differences compared to canonical H3. A loop region in CENP-A protrudes from the nucleosome, exposing residues arginine 80 and glycine 81 (RG-loop) and thereby creating a site for CENP-N and CENP-C recruitment (**Figure 1A,B**). A second key difference is that the αN helix of CENP-A is one turn shorter than in canonical H3, reducing DNA contacts and thus increasing the flexibility of DNA at the nucleosome entry and exit sites^15–19^ (**Figure 1A,B**). In accordance, recent cryogenic electron microscopy (cryo-EM) structures show that yeast CENP-A/Cse4 and human CENP-A nucleosomes form open structures when bound to core kinetochore components^20,21^ as well as induce untwisting of trinucleosomes^19^ when flanked by canonical H3-containing nucleosomes. This flexibility has been shown to be essential for proper chromosome segregation and the recruitment of CENP-B and CENP-C^16^. However, the question of how the increased flexibility of CENP-A nucleosomes alters chromatin structure remains unclear.

**Figure 1:**
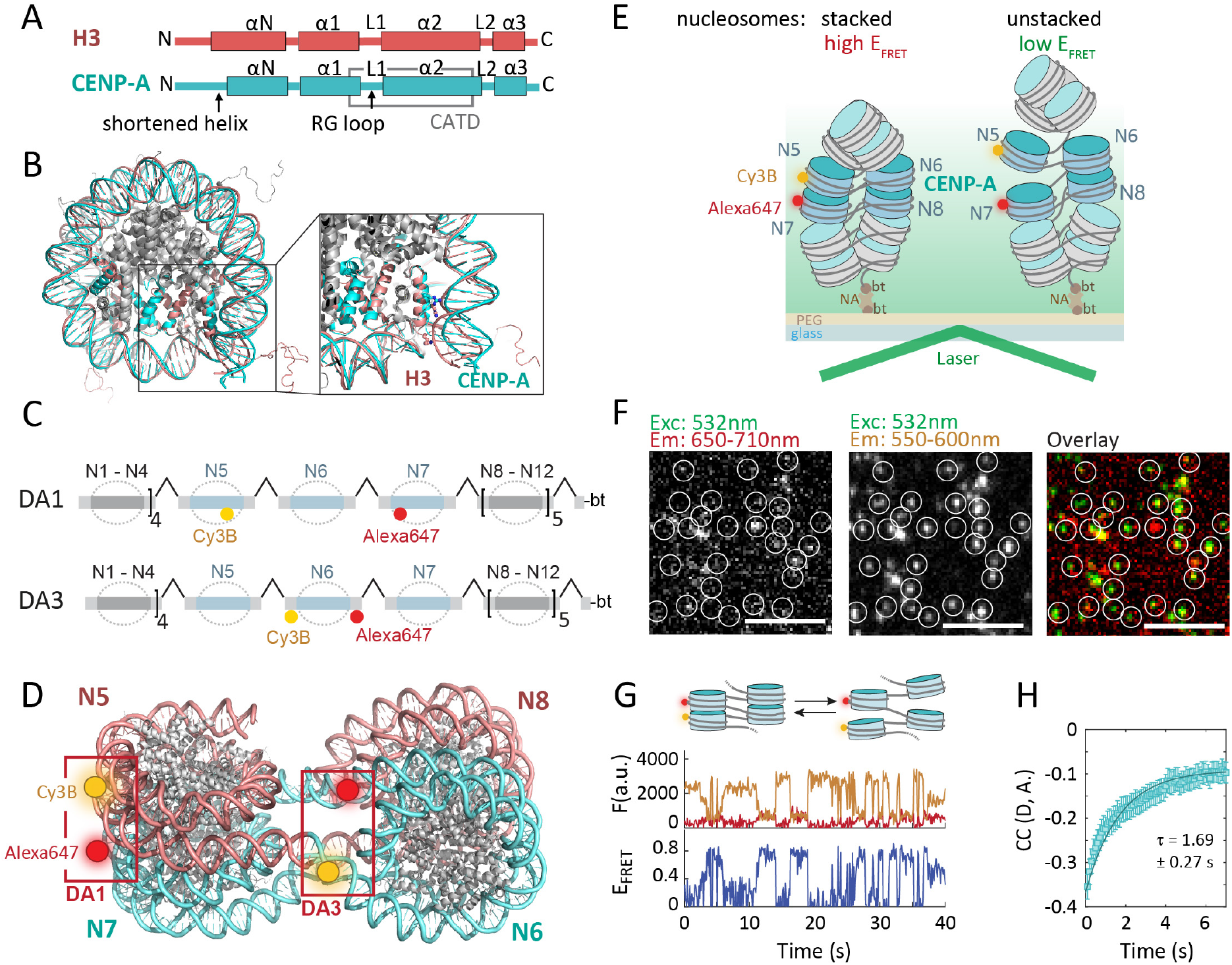
CENP-A has a dynamic chromatin structure. **A**) Comparison of the sequence and secondary structure organization of H3 and CENP-A. L1,2: loop 1,2; α1-3: helical segments 1-3; CATD: CENP-A targeting domain. **B**) Differences in H3 and CENP-A nucleosomes, showing reduced DNA contacts of CENP-A resulting in more dynamic DNA. (PDB code: 1KX5 (H3, red), 6SE0 (CENP-A, cyan)). **C**) Scheme of chromatin DNA assembly containing FRET donor (Cy3B, yellow) and acceptor (Alexa Fluor 647, [Alexa647], red) at nucleosome N5 and N7 (DA1, top) and at N6 (DA3, bottom). **D**) Tetranucleosome structure (PDB code: 1ZBB) showing stacked nucleosomes in a two-start organization. DA1 and DA3 FRET dye pairs are indicated. **E**) Scheme of smFRET – TIRF experiment (DA1 illustrated). **F**) Microscopic images showing FRET data of single CENP-A-containing chromatin arrays at 4 mM Mg^2+^, white circles indicate the positions of immobilized fluorescently tagged chromatin, scale bar: 5 μm. **G**) Fluorescence and FRET traces of CENP-A chromatin fibers, carrying DA1 labels at 4 mM Mg^2+^, showing donor dye (orange), acceptor dye (dark red) emissions, and calculated E_FRET_ trace (blue). **H**) Donor–acceptor channel cross-correlation analysis of DA1-labeled CENP-A chromatin fibers. Fit: cross-correlation relaxation time *τ* = 1.69 ±0.27 s.

Apart from the centromeric histone variant CENP-A, the constitutive centromeric protein CENP-B is the only kinetochore component to make direct and specific DNA contacts^22^. It recognizes a defined DNA sequence, the CENP-B box motif (B-Box) within alpha-satellite repeats^23–25^ via its N-terminal DNA-binding domain. While CENP-B is nonessential for centromere function^26–28^, it has been shown to be required for *de novo* centromere assembly in human artificial chromosomes (HACs)^29–31^, as well as to stabilize the kinetochore and enhance centromere function^32,33^. Indeed, depletion of CENP-B greatly increases cell lethality upon CENP-A loss^34^. Finally, CENP-B acts as an anchor for diverse chromatin proteins, including heterochromatin protein 1 (HP1) and the H3K36-specific methyltransferase ASH1L^35^. These combined functions render CENP-B a key regulator of the centromeric chromatin state, and consequently, an important factor controlling the structure and stability of the centromere.

B-Boxes are regularly distributed in alpha-satellite DNA, occurring generally within every second repeat and being localized near the DNA entry-exit region of centromeric nucleosomes^36^. Binding these DNA sequences requires CENP-B to invade centromeric chromatin structure. CENP-B localization is not limited to CENP-A nucleosomes^23^; however, previous studies implied that its DNA-binding domain (DBD) can make preferential contacts to CENP-A through interaction with the amino tail of CENP-A^32,33^. Moreover, CENP-B is able to homodimerize through its C-terminal domain and bridge B-Boxes in two alpha-satellite monomers^37^, forming chromatin loops^38^. The possibility for CENP-B to invade centro-chromatin structure and to cross-bridge chromatin elements requires significant flexibility of centromeric chromatin. How CENP-B remodels centromeric chromatin structure is, however, an unsolved question.

We have recently developed single-molecule fluorescence resonance energy transfer (smFRET) methods to reveal dynamic chromatin organization^39^. These studies revealed that chromatin exhibits micro-to millisecond conformational fluctuations, which transiently expose internal DNA segments. Such structural fluctuations are exploited by DNA-binding proteins, e.g. transcription factors, to access their binding sites within compact chromatin^40^. Here, we employ single-molecule fluorescence methods, including smFRET and single-molecule colocalization imaging, to probe the dynamic structure of centro-chromatin. Using reconstituted CENP-A chromatin, we find that centromeric histones induce a highly dynamic chromatin state, which enables unhindered DNA access to CENP-B. CENP-B binding induces further chromatin opening by distorting the bound linker DNA containing its B-Box binding sequences. Finally, we find that the two centromeric proteins dynamically collaborate in cells, where the presence of CENP-A stabilizes CENP-B retention at centromeres. In conclusion, we demonstrate a chromatin-mediated crosstalk between two key centromeric proteins, establishing a dynamic and accessible chromatin state at centromeres.

## Results

### CENP-A creates a dynamic chromatin structure

Multiple structural studies have indicated that CENP-A mononucleosomes are more flexible compared to H3 nucleosomes^15,19,41,42^, although how this increased local flexibility modulates longer-range higher-order chromatin structure is unclear. Increased nucleosomal flexibility might lead to a more dynamic higher-order structure. Indeed, cryo-EM analysis of CENP-A containing tri-nucleosomes suggested a highly heterogeneous untwisted structure^19^. Conversely, CENP-A might allow for the formation of compact structures by relieving conformational strain. In accordance with this hypothesis, sedimentation assays previously showed that CENP-A increased the formation of compacted chromatin structures^41,43^.

We decided to probe the dynamic organization of CENP-A-containing centro-chromatin by employing a recently established single-molecule fluorescence resonance energy transfer (smFRET) method, measuring energy transfer between two fluorescent dyes placed at defined positions within chromatin fibers and reporting on chromatin structural parameters^39,44^. Using this approach and varying FRET donor and acceptor dye placement, we can directly detect nucleosome stacking or determine the orientation of linker DNA on a singlefiber level. To implement the approach, we reconstituted 12-mer chromatin fibers using histone octamers containing either CENP-A or H3, as well as all other canonical human histone proteins, namely H4, H2A and H2B. The DNA template used for chromatin fiber assembly contained 12 tandem repeats of the 601 nucleosome positioning sequence (NPS)^45^, ensuring nucleosome placement with base pair accuracy, which is instrumental for smFRET measurements. Individual 601 NPSs were further separated by 30 bp linker DNA (**Figure 1C**). FRET donor (Cy3B) and acceptor dyes (Alexa647) were placed on nucleosome 5 (N5) and N7 (called the donor-acceptor 1 (DA1) pair^39^, **Figure 1D**) using a DNA plug-and-play strategy^46^ (see **Figure S1** for analytical data of reconstitutions and **Tables S1-3** for all DNA constructs). This FRET pair allows the detection of nucleosome stacking, i.e. face-to-face contact between next-neighbor nucleosomes (**Fig. 1D**). To probe the relative orientation of linker DNA, we further used alternative FRET dye positions, named DA3^39^, where the dyes flank the central nucleosome N6 (**Figure 1C,D**, **Figure S1**, **Tables S1-3**). Together, these two labeling schemes enable us to probe both DNA orientation and nucleosome stacking within the chromatin fiber.

Reconstituted CENP-A- or H3-containing chromatin fibers, hereon called CENP-A chromatin or H3 chromatin, respectively, were then immobilized in a flow channel, followed by single-molecule fluorescence imaging via total internal reflection fluorescence (smTIRF) microscopy (**Figure 1E**). Individual chromatin fibers were detected as diffraction-limited spots in the donor- and acceptor-emission channels (**Figure 1F**). We recorded donor and acceptor dye fluorescence emission over time, and calculated time-traces of FRET efficiency (E_FRET_) (**Figure 1G**, **Methods**). For H3 chromatin fibers at 4 mM Mg^2+^, conditions that induce chromatin compaction via shielding of negative DNA charges^47^, this resulted in stable E_FRET_ ∼ 0.5 over several tens of seconds^39^. This is in agreement with previous experiments that used confocal single-molecule fluorescence spectroscopy of H3-containing chromatin to reveal rapid structural fluctuations on the μs – ms timescale^39^, which are mostly undetectable on the slower smTIRF timescale. In contrast, when observing CENP-A chromatin under the same conditions, we observed large-scale fluctuations in FRET efficiency on the seconds timescale for a substantial fraction of CENP-A-containing chromatin fibers (**Figure 1G**). Donor-acceptor cross-correlation analysis of these fluctuations, including all traces that showed an average E_FRET_ > 0.2, revealed large-amplitude motions with a relaxation time τ_R_ = 1.69 ± 0.27 s (**Figure 1H**). In comparison, H3 chromatin showed only small-amplitude motions with τ_R_ ∼ 0.1 s when analyzed similarly^39^. Together, these results show that CENP-A renders chromatin fibers to be much more dynamic, as opposed to more stably folded H3 chromatin.

### CENP-A disrupts nucleosome-stacking interactions

Comparing the distribution of E_FRET_ values between H3 and CENP-A chromatin provided further insights into the organization of centromeric chromatin. We constructed histograms of E_FRET_ values, integrating all time traces. The resulting distributions could be described by the sum of two Gaussian distributions, corresponding to a low E_FRET_ (open) and mid-to-high E_FRET_ (compacted) conformation (**Fig. 2A**, **Table S4**).

**Figure 2.**
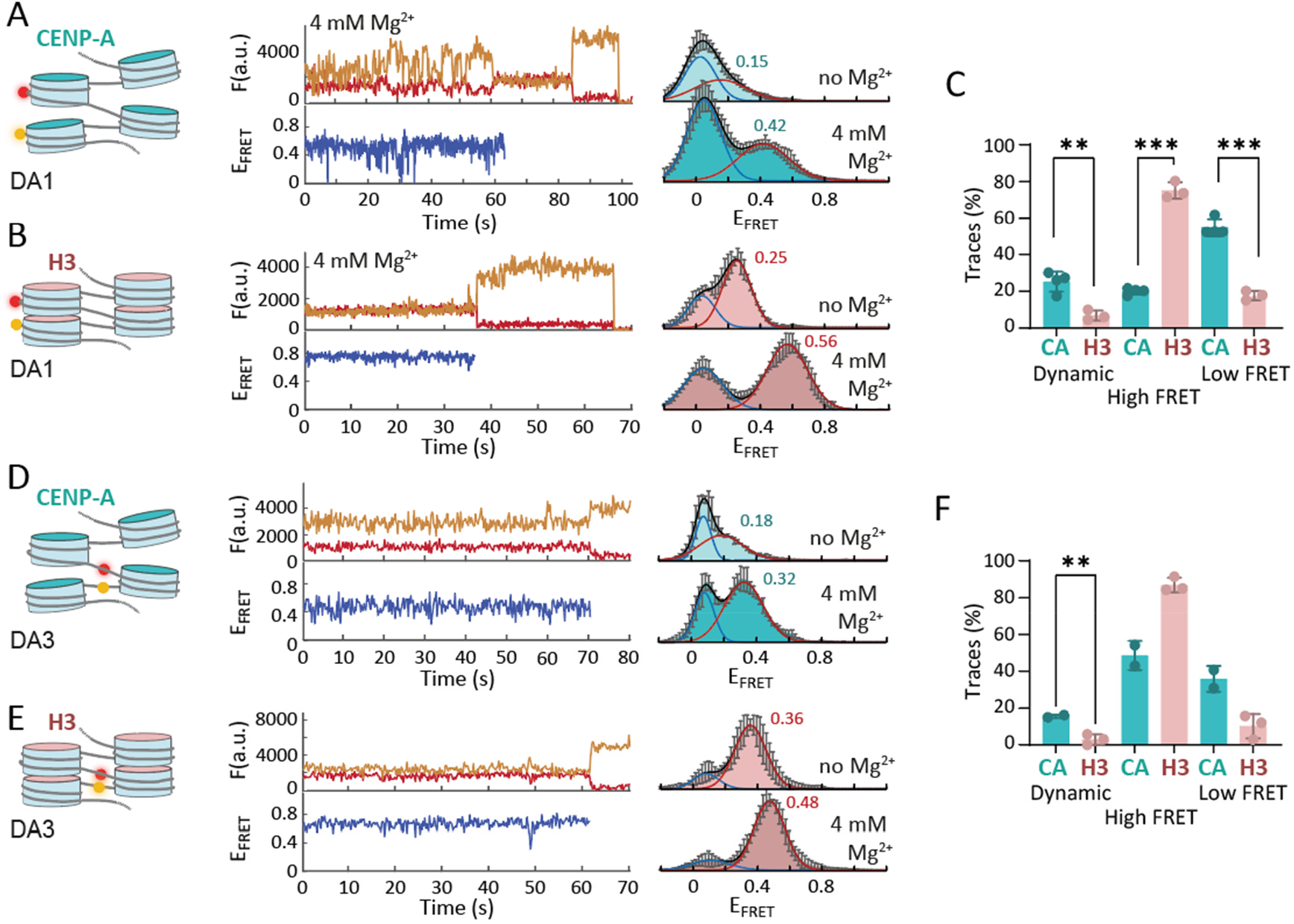
CENP-A generates open chromatin. **A**) Left: Single-molecule traces (donor: orange, acceptor: red, FRET: blue) for CENP-A chromatin, labeled at DA1, at 0 mM Mg^2+^ (top), 4 mM Mg^2+^ (bottom) until either donor or acceptor dye were photobleached. Right: FRET population histograms, observed for CENP-A chromatin, labeled at DA1, at the indicated Mg^2+^ concentrations. All Histograms are the average of n>2 independent repeats. Histograms are fitted with Gaussian functions (red). For all fit values, see **Table S4**. Peak E_FRET_ for high FRET populations are indicated. Error bars are standard error. **B**) Single-molecule traces for H3 chromatin, labeled at DA1, and accompanying FRET histograms at the indicated Mg^2+^ concentrations. **C**) Percentage of dynamic, high-FRET, and low-FRET traces of CENP-A (CA) and H3 chromatin for DA1. Error bars show S.D. \**P*<0.05, \*\**P*<0.01,\*\*\**P*<0.001 using 2-tailed unpaired t-test. **D**) Single-molecule traces for CENP-A chromatin, labeled at DA3, and accompanying FRET histograms at the indicated Mg^2+^ concentrations. **E**) Single-molecule traces for H3 chromatin, labeled at DA3, and accompanying FRET histograms at the indicated Mg^2+^ concentrations. **F**) Percentage of dynamic, high-FRET, and low-FRET traces of CENP-A (CA) and H3 chromatin for DA3. Statistics are calculated as in **C**).

First, we focused on chromatin fibers containing the DA1 dye pair, monitoring stacking interactions between next-neighbor nucleosomes (N5-N7). At low ionic strength (40 mM KCl), we observed that CENP-A chromatin was essentially open with a majority of molecules showing low FRET (E_FRET_ < 0.1, **Figure 2A**). H3 chromatin, in contrast, exhibited a majority population of intermediate FRET efficiency (E_FRET_ = 0.2 - 0.5, **Figure 2B**). This is consistent with the existence of rapidly exchanging populations of open or stacked nucleosomes, with exchange kinetics on the μs – ms timescale^39^. Addition of bivalent cations (4 mM Mg^2+^) induced chromatin compaction in both CENP-A and H3 chromatin, as exhibited by a high-FRET population centered at E_FRET_ values of 0.44 for CENP-A and 0.54 for H3 chromatin (**Figure 2A,B**). This population was markedly smaller for CENP-A, and characterized by a broad distribution of E_FRET_ values due to large-scale conformational dynamics in the seconds timescale (**Figures 2A** and **1G**). We then quantified the number of single-molecule traces, which (i) showed large-scale fluctuations as in **Figure 2A** (labeled as ‘Dynamic’), (ii) showed stable medium-to-high E_FRET_ values (labeled as ‘High FRET’) or (iii) exhibited low E_FRET_ values (labeled as ‘Low FRET’) for both CENP-A and H3 chromatin (**Figure 2C**). Also, in this analysis, CENP-A chromatin showed a much lower percentage of traces at high E_FRET_ compared to H3 chromatin, but a significantly increased percentage of dynamic traces, corroborating its open and dynamic organization.

Alternatively, detecting FRET via the DA3 FRET pair provides information on the orientation of the linker DNA that flanks the central nucleosome within the reconstituted fibers (N6, **Figure 1D**). In CENP-A chromatin, we observed two populations of low- and mid-FRET (E_FRET_ = 0.18) values at low ionic strength, indicating that the DNA in CENP-A nucleosomes is divergent at the entry-exit sites, even in a chromatin fiber context. Upon the addition of 4 mM Mg^2+^, a higher FRET state was populated (E_FRET_ = 0.32), indicating transient higher-order structure stabilization. Conversely, the distance between DNA strands at the nucleosome entry-exit sites was smaller in H3 chromatin (**Figure 2E**), as observed by higher FRET values for both low ionic strength (E_FRET_ = 0.36) and at 4 mM Mg^2+^ (E_FRET_ = 0.48). From the DA3 vantage point which monitors the relative position of the linker DNA, dynamics were less pronounced but still more prevalent in CENP-A chromatin compared to H3 chromatin (**Figure 2F**). To gain further insights into the conformation of CENP-A chromatin, we compared measured E_FRET_ to calculated E_FRET_ values that were generated using cryo-EM structures of H3-H3-H3 or H3-CENP-A-H3 trinucleosomes^19^ (**Figure S2**). Our measurements for H3 were compatible with expected values from the cryo-EM structure. In contrast, our measured E_FRET_ values were higher than those expected from the structure of a H3-CENP-A-H3 trinucleosome exhibiting an untwisted central nucleosome. Our results thus indicate that fiber interactions constrain CENP-A nucleosomes into a more H3-like conformation in an extended fiber context (**Figure S2**).

Together, our data show that CENP-A nucleosomes can engage in stacking interactions, forming tetranucleosome units, albeit at lower efficiency and in a distorted architecture. Moreover, CENP-A chromatin exhibits greatly increased dynamics associated with large-scale breathing motions on the seconds timescale. Both the distorted structure and dynamic openings may result in increased local DNA access within centromeric regions.

### CENP-B efficiently binds nucleosomes independent of histone type

We thus wondered if CENP-A chromatin incorporation may provide a particularly permissive chromatin state for DNA-binding proteins. CENP-B is a DNA-binding factor of particular relevance, as it marks centromeres and is instrumental for the establishment of a centro-chromatin state^38^. To determine how CENP-A controls DNA access for CENP-B, we directly determined the binding dynamics of full-length CENP-B using single-molecule colocalization imaging^40,48^ (**Figure 3A**). In such an experiment, individual naked DNA molecules, reconstituted nucleosomes or chromatin fibers, all labeled with a far-red fluorescent dye are surface-immobilized within a microfluidic flow-channel (**Figure 3A**). Their individual positions are then detected via single-molecule total internal reflection fluorescence (smTIRF) microscopy. Subsequently, recombinant full-length CENP-B, which is labeled by a fluorescent dye in the green-orange spectrum is injected into the channel, and individual dynamic binding events are detected by colocalization imaging between the far-red (DNA/chromatin) and orange (CENP-B) channels (**Figure 3A**).

**Figure 3:**
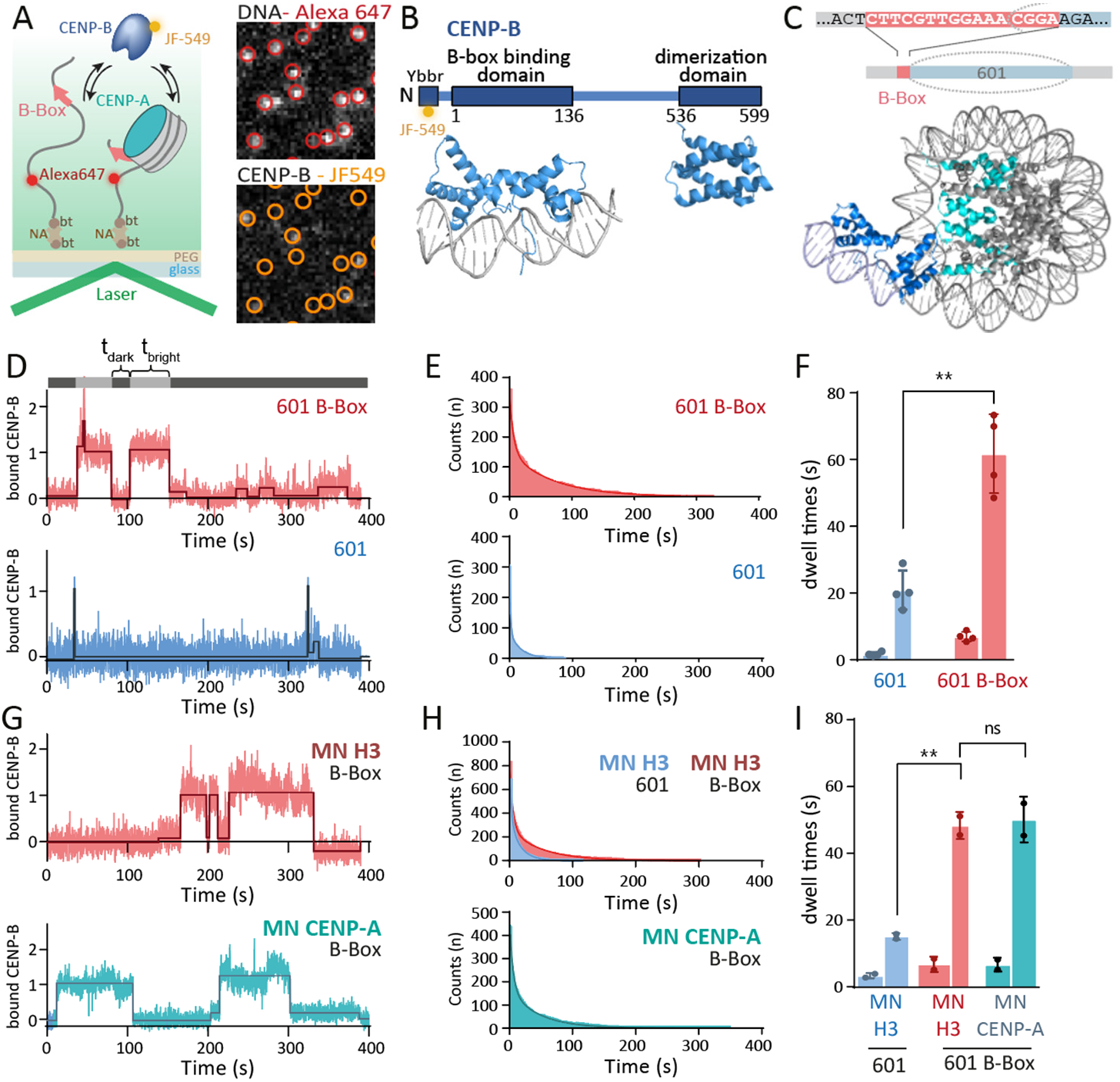
CENP-B binds independently of histone type. **A**) Left: Scheme of smTIRF experiment to detect CENP-B binding to B-Box- or non-B-Box-containing 601 DNA or nucleosomes. bt: biotin, NA: neutravidin. Right: smTIRF image showing immobilized 601-B-Box DNA in the far red channel (top, red circles) and CENP-B binding in the green-orange channel (bottom). **B**) Used CENP-B construct. Ybbr: peptide tag for fluorescent labeling. Below: Structures of the DNA-binding domain (PDB code: 1HLV) and dimerization domain (PDB code: 1UFI). **C**) Top: Position of the B-Box DNA sequence relative to the CENP-A nucleosome. Bottom: Model of a CENP-A nucleosome containing a B-Box-bound CENP-B. **D**) Fluorescence time trace (blue) of CENP-B binding events to 601-B-Box (top) and non-B-Box DNA (bottom). Free (t_dark_) and bound times (t_bright_) are determined via thresholding. **E**) Cumulative histogram of CENP-B binding to B-Box- (red) and non-B-Box- (blue) containing DNA, fitted by a bi-exponential function (solid line). **F**) Dissociation time constants (*τ_off,1_* and *τ_off,2_*) of CENP-B to non-B-Box (blue) and B-Box (red) 601 DNA. Width of bars indicates relative amplitudes A_1_ and A_2_. For all fit parameters, see **Table S5**. n=4; error bars are S.D. *\*\*P*< 0.01, 2-tailed unpaired t-test. **G**) Representative fluorescence time trace (blue) of CENP-B binding events to 601-B-Box nucleosomes (MN) containing H3 (top) or CENP-A (bottom). **H**) Cumulative histogram of CENP-B binding to B-Box nucleosomes, containing H3 (top) or CENP-A (bottom) fitted by a bi-exponential function (solid line). **I**) Dissociation time constants of CENP-B to H3 non-B-Box (blue), H3 B-Box (red) and CENP-A B-Box (grey) 601 nucleosomes. n=2; error bars: S.D; \*\**P*< 0.01, ns: *P* > 0.05, 2-tailed unpaired t-test.

To implement this experiment, we prepared recombinant full-length CENP-B using a DNA construct that contained both the CENP-B DNA-binding and dimerization domains and carrying an N-terminal peptide tag that allowed chemoenzymatic labeling^49^ with a high-performance green-orange dye (JF-549^50^, **Figure 3B** and **Figure S3**). Overall labeling efficiency was around 50%, ensuring that most CENP-B dimers carried one fluorophore. In parallel, we generated a DNA template composed of a 601 NPS, either without any sequence changes (601) or containing a CENP-B Box (601 B-Box) that was positioned 69 bp from the 601 dyad position, similar to previous studies^33^. This position mimics the location of the B-Box relative to nucleosomes within human alphoid DNA^36^ (**Figure 3C**, **Figure S3** and **Tables S1-3**). The DNA was further modified with a far-red fluorescent dye (Alexa Fluor 647) and a biotin moiety for immobilization.

We proceeded to measure CENP-B binding dynamics to immobilized naked DNA, with or without a B-Box, via single-molecule colocalization imaging. For these experiments, we injected CENP-B at a concentration chosen such that we were able to clearly observe single CENP-B binding events (approx. 5 nM) and under native ionic strength (150 mM salt) (**Figure 3A,D**). For each detected DNA molecule in the far-red channel, we then generated time traces of CENP-B binding observed in the green-orange channel, which allowed us to determine the length of individual binding events (*t_bright_*) and the time between binding interactions (*t_dark_*, **Figure 3D**). Lifetime histograms of binding events, generated from *t_bright_*, provided information about CENP-B residence times, whereas histograms of *t_dark_* reported on binding rates (**Figure S4**). Focusing on residence times, we analyzed the lifetime histograms of the bound state using a bi-exponential fit, resulting in a short (*τ*_off,1_) and long (*τ*_off,2_) residence times (**Figure 3E**, for all fit parameters, see **Table S5**). This is commonly observed for chromatin binding proteins^40,48,51^, and indicates different interaction modes, usually interpreted as non-specific charge-based interactions (*τ*_off,1_) and engagement of specific DNA sequences via the DBD (*τ*_off,2_). In the presence of a B-Box, around half of all CENP-B-DNA interaction exhibited a short residence time (*τ*_off,1_ = 6.9 ± 1.4 s), whereas the remainder showed a long residence time (*τ*_off,2_ = 61 ± 11 s). This is consistent with a mid-nanomolar affinity, as determined by electrophoretic mobility shift assays (**Figure S5**), and also with previous results^38^. These long, multi-second residence times, comparable to those of transcription factors^40,52,53^, enable CENP-B to play a role in chromatin structure organization, while ensuring a dynamic chromatin state. Importantly, full-length CENP-B was required for stable DNA binding: residence times for a short CENP-B construct, encompassing only the DNA-binding domain (residues 1 – 150), were strongly reduced (*τ*_off,1_ = 2.6 ± 0.2 s, *τ*_off,2_ = 22.8 ± 4.4 s) (**Figure S6**).

These findings suggest that the kinetochore-binding and dimerization domains enhance B-Box recognition by CENP-B. Finally, the absence of a B-Box sequence resulted in significantly shorter binding times for full-length CENP-B (*τ*_off,1_ = 1.6 ± 0.6 s, *τ*_off,2_ = 20 ± 6 s). In conclusion, CENP-B can also bind DNA in a sequence-independent fashion, possibly allowing an efficient search process for B-Box sequence elements in the genome^54^.

We then tested whether CENP-B can efficiently invade and bind to nucleosomal DNA. To this end, we reconstituted either H3- or CENP-A-containing nucleosomes on the previously used DNA templates (**Figure S3**), with the CENP-B box precisely positioned at the DNA-entry-exit site as in human alphoid DNA^36^. These nucleosomes were used to measure CENP-B binding dynamics via single-molecule colocalization imaging (**Figure 3G**). Interestingly, the presence of H3-containing histone octamers did not greatly affect the observed CENP-B residence times (**Figures 3G-I**). CENP-B was efficiently able to invade (**Figure S4**) and remain bound at H3-containing mononucleosomes with residence times *τ*_off,1_ = 6.9 ± 2.3 s (66% of all events) and *τ*_off,2_ = 48 ± 4 s (34% of all events), not significantly different compared to those for naked DNA (**Figures 3I**).

Similarly, CENP-B was able to bind CENP-A-containing mononucleosomes basically unhindered, with residence times *τ*_off,1_ = 6.8 ± 2.06 s (51% of all events) and *τ*_off,2_ = 50 ± 7 s (49% of all events, **Figures 3G-I**, see **Table S5** for all data). Within CENP-A nucleosomes, the peripheral DNA is accessible^15^ and thus the CENP-B box is exposed, which explains a negligible effect on CENP-B binding kinetics. Conversely, we also could not detect any positive effect of CENP-A to promote the binding of full-length CENP-B, e.g. via direct interactions as previously suggested from experiments using the CENP-B DBD alone^33^. In H3-containing nucleosomes, the CENP-B box is partially located within the nucleosome structure (**Figure 3C**). The fact that nucleosomes do not reduce CENP-B binding or residence times in this context therefore indicates that the interactions of CENP-B to its 17-bp DNA recognition sequence can efficiently outcompete peripheral histone-DNA contacts, independently of histone type.

### CENP-B binding is affected by the underlying chromatin state

In the cell, CENP-B does not interact with individual nucleosomes but binds to extended chromatin fibers, which can adopt various conformations and higher-order structures^1^. While our single-molecule experiments revealed that CENP-B binding is not enhanced by CENP-A in a single-nucleosome context, we hypothesized that the open and dynamic nature of centromeric chromatin may present a preferred binding environment.

We proceeded to test this hypothesis by measuring CENP-B binding kinetics to reconstituted 12-mer chromatin fibers containing either H3 or CENP-A (**Figure 4A**). Here, we positioned a single B-Box site within the central nucleosome (N6) at a distance of 69 bp from the dyad (**Figure 4B**), similar to our previous binding studies with mononucleosomes (**Figure 3**). These chromatin fibers containing either H3 or CENP-A were then immobilized in flow cells to determine CENP-B binding kinetics by single-molecule colocalization imaging. Importantly, we performed these experiments under physiological salt conditions (150 mM KCl) that result in a compacted chromatin structure^40^.

**Figure 4:**
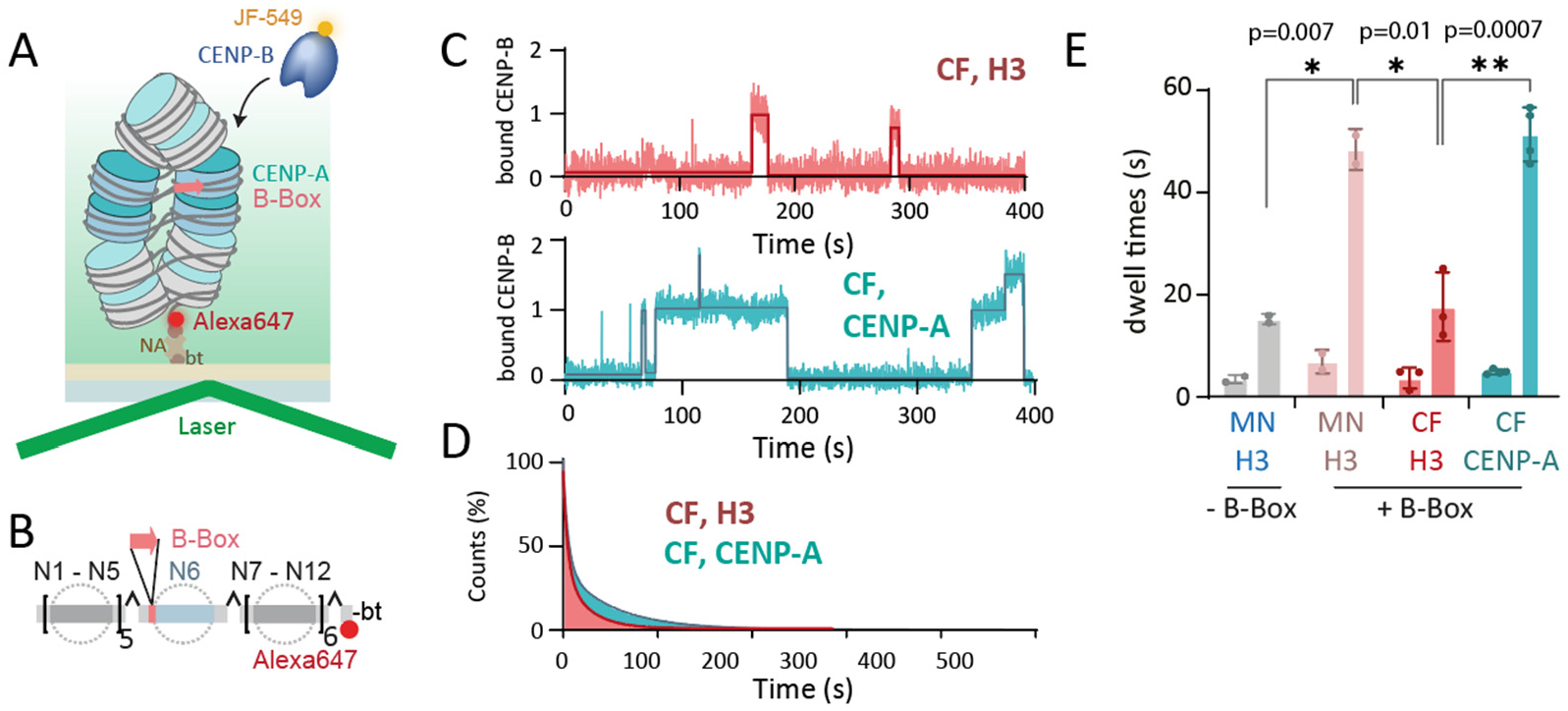
CENP-B binding is controlled by the local chromatin state. **A**) Scheme of colocalization experiment to detect CENP-B binding to immobilized H3- or CENP-A-containing chromatin fibers. **B**) Scheme of array DNA assembly containing one B-Box at nucleosome N6 (bottom). bt is biotin, NA is neutravidin. **C**) Representative fluorescence time trace of CENP-B binding events to H3- (top) and CENP-A- (bottom) containing chromatin fibers (CF). The traces are fitted by a thresholding algorithm. **D**) Cumulative histogram of CENP-B binding to H3 arrays (orange) and CENP-A arrays (green) fitted by a bi-exponential function (solid line). **E**) Dissociation time constants (*τ_off,1_* and *τ_off,2_*) of CENP-B to the indicated nucleosome and chromatin constructs. CF: chromatin fiber. Thickness of the bars indicate amplitudes A_i_ associated with the indicated dwell times. n=2-4; error bars are S.D.

In H3 chromatin, the detected specific residence times *τ*_off,2_ for CENP-B were substantially reduced by almost 3-fold to *τ*_off,2_ = 17.6 ± 6.7 s compared to those in mononucleosomes (**Figure 4C-E**). X-ray crystallography studies revealed that B-Box DNA is distorted in the CENP-B-bound state^25,55^. Such distortion of the linker DNA is most likely incompatible with stacked nucleosome conformations that are observed in structures of compact tetranucleosomes or chromatin fibers^19,56,57^. Thus, CENP-B may not be able to form an optimal DNA-bound complex, resulting in its rapid eviction from chromatin fiber structure.

Conversely, we did not observe such inhibition in chromatin fibers containing CENP-A nucleosomes: Here, specific residence times for CENP-B were restored to *τ*_off,2_ = 51.4 ± 5.3 s, similar to those in free mononucleosomes (**Fig 4C-E**). This recovery of binding lifetimes within CENP-A-containing chromatin suggest that CENP-A chromatin fibers can accommodate CENP-B, tolerating bent DNA, and provide less hindered DNA access.

To corroborate this hypothesis, we measured the conformation of the linker DNA using smFRET, using linker-positioned dyes (DA3). We indeed detected a loss of E_FRET_ in CENP-A chromatin upon CENP-B binding, indicating that linker DNA is pried open (**Figure S7**). Again, the effect was not detectable in H3 chromatin, where DNA is more rigidly held in place by histone contacts (**Figure S7**). In conclusion, CENP-A generates a permissible environment for chromatin invasion of DNA-binding factors, due to its effect of increasing local chromatin flexibility.

### CENP-B binding pries opens chromatin structure

A critical function of CENP-B is its ability to cross-bridge chromatin elements and induce loops in chromatin structure, as it exists as a dimer able to bind to B-Box DNA motifs simultaneously^38^. We thus wondered if CENP-B would be able to stabilize next-neighbor nucleosome interactions within chromatin structure (**Figure 5A**). We reconstituted H3 or CENP-A chromatin fibers on a DNA template either with or without B-Boxes positioned in nucleosomes N5 and N7, as well as containing fluorescent labels to detect nucleosome stacking (DA1, **Figure 5B** and **Figure S1**). This arrangement of B-Boxes is similar to their occurrence in native human centromeres^36^. We then proceeded to titrate CENP-B to the reconstituted chromatin fibers and detected structural changes via smFRET. H3 chromatin showed a high-FRET population (E_FRET_ = 0.56) in the absence of CENP-B (**Figure 5C**, left and middle, **Table S6**). Adding CENP-B up to a concentration of 50 nM had a negligible effect on H3 chromatin fibers lacking B-Boxes (**Figure 5C, D**). However, when exposing H3 chromatin containing two B-Boxes in next-neighbor nucleosomes to increasing CENP-B concentrations, we observed a decrease in the high-FRET population and a simultaneous increase in the low-FRET population (**Figure 5C, E**). Transient CENP-B binding thus results in a substantial loss of nucleosome stacking, even though the linker DNA orientation in H3 chromatin is not strongly altered under the same conditions (**Figure S7**). When performing these experiments with CENP-A chromatin fibers, which already exist in a dynamic and open state, we also observed a depletion of the high-FRET population, resulting in an almost complete loss of detectable FRET at 50 nM CENP-B (**Figure 5C, F**).

**Figure 5:**
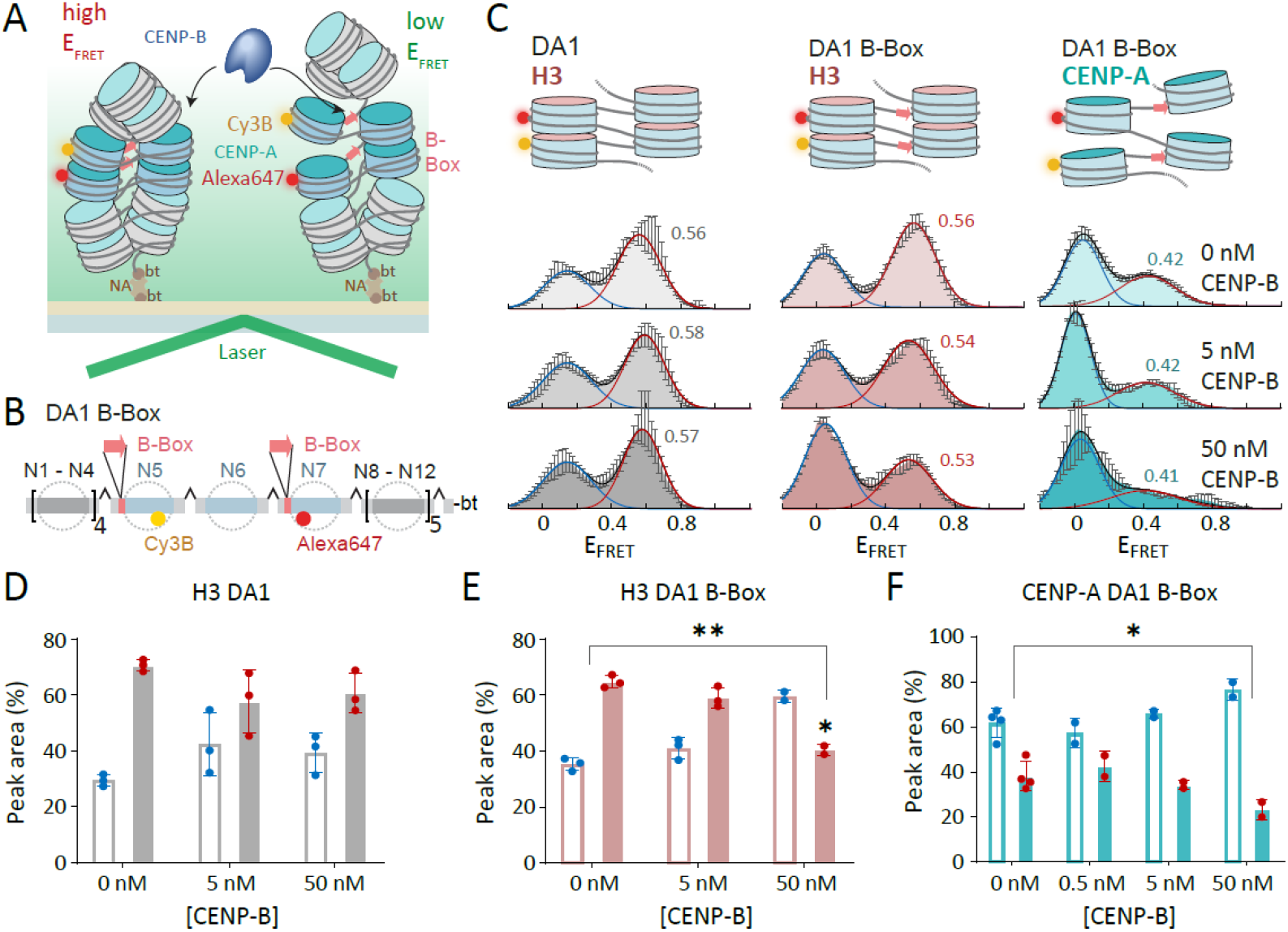
CENP-B can induce local unstacking of chromatin. **A**) Scheme of smFRET experiment. **B**) Scheme of chromatin DNA containing B-Box sites at nucleosome N5 and N7 with FRET donor (Cy3B, yellow) and acceptor (Alexa Fluor 647, red) dyes at nucleosome N5 and N7 (DA1BB, top). **C**) Illustrative structures (left) and histograms of E_FRET_ of chromatin fibers at 4mM Mg^2+^ concentration (right). Rows indicate histograms for H3 DA1, H3 DA1BB and CENP-A DA3BB, respectively. Columns indicate CENP-B concentration. Histograms are the average of n=2-4 independent repeats. Histograms are fitted with Gaussian functions (red). Error bars are standard error. **D**) Percentage of low- (blue) and high-FRET (red) sub-populations for H3-containing chromatin arrays lacking B-Boxes (DA1) at the indicated CENP-B concentrations. Error bars show S.D. **E**) Percentage of low- (blue) and high-FRET (red) sub-populations for H3-containing chromatin arrays with B-Boxes (DA1BB) at the indicated CENP-B concentrations. Error bars show S.D. \*\**P*<0.01 for 0 nM CENP-B vs 50 nM CENP-B, \**P*<0.05 for 50 nM CENP-B H3 DA1 vs H3 DA1BB using 2-tailed unpaired *t*-test. **F**) Percentage of low- (blue) and high-FRET (red) sub-populations for CENP-A-containing chromatin arrays with B-Boxes (DA1BB) at the indicated CENP-B concentrations. Error bars show S.D. \**P*<0.05 for 0 nM CENP-B vs 50 nM CENP-B using 2-tailed unpaired *t*-test.

In conclusion, we did not observe the stabilization of nucleosome interactions by chromatin-bound CENP-B. In contrast, our measurements indicate that CENP-B induces a more open chromatin state, which is characterized by larger inter-nucleosome distances. This may involve the formation of local loop-like structures or of alternative nucleosome interactions leading to larger inter-dye distances, or may just be a consequence of steric clashes of bound CENP-B with neighboring nucleosomes, as observed for transcription factors^40^.

### CENP-A stabilizes CENP-B binding in cells

Taken together, our results suggest CENP-B recruitment to the centromere is affected by the underlying chromatin state, with the more open CENP-A-containing chromatin more easily able to recruit CENP-B. Simultaneously, CENP-B, when recruited to the B-Box, is able to induce local opening of chromatin and stabilize alternative chromatin structures. Based on these results, we wondered whether CENP-B-chromatin interactions are also stabilized by CENP-A in living cells. We thus determined CENP-B dynamics in the presence and absence of CENP-A via fluorescence recovery after photobleaching (FRAP). A fluorescent protein (mEos3.2) was fused to CENP-B, placed under a doxycycline-inducible promoter and stably integrated into DLD-1 cells (human adenocarcinoma). Upon induction with doxycycline, CENP-B was expressed and localized to centromeres as judged by immunofluorescence imaging using CENP-C as a centromere marker (**Figure S8**). We then bleached two kinetochores per cell for 20 – 30 cells and determined the kinetics of CENP-B recovery in the presence of CENP-A (**Figure 6A**). Analyzing the averaged recovery traces revealed a time constant of fluorescence recovery, *τ*_FRAP_ = 220 s, and an immobile fraction of 38% (**Figure 6B**). This is consistent with previously reported recovery rates for CENP-B^58^. We then depleted CENP-A via short interfering RNA (siRNA) treatment for 48 h and confirmed the strong reduction of CENP-A levels in cells via immunoblotting (**Figure S8, Table S7**). Subsequently, we performed FRAP of fluorescently labeled CENP-B in these CENP-A-depleted cells (**Figure 6A**) and, indeed, observed a faster recovery with *τ*_FRAP_ = 136 s and a reduced immobile protein fraction of 26%, indicating a weakening of CENP-B binding to chromatin (**Figure 6B**). Importantly, this effect was only observed for siRNA against CENP-B and not for scrambled control siRNA (**Figure S9**). We further assessed the effect of CENP-A depletion for a truncated version of CENP-B. The recovery of the construct CENP-B_1-500_, which lacks the dimerization domain, was still sensitive to CENP-A levels, albeit the effect of depleting CENP-A was weaker (**Figure S10**). Conversely, CENP-B_1-150_ was highly dynamic and its binding to chromatin was mostly unaffected by CENP-A levels (**Figure S10**).

**Figure 6:**
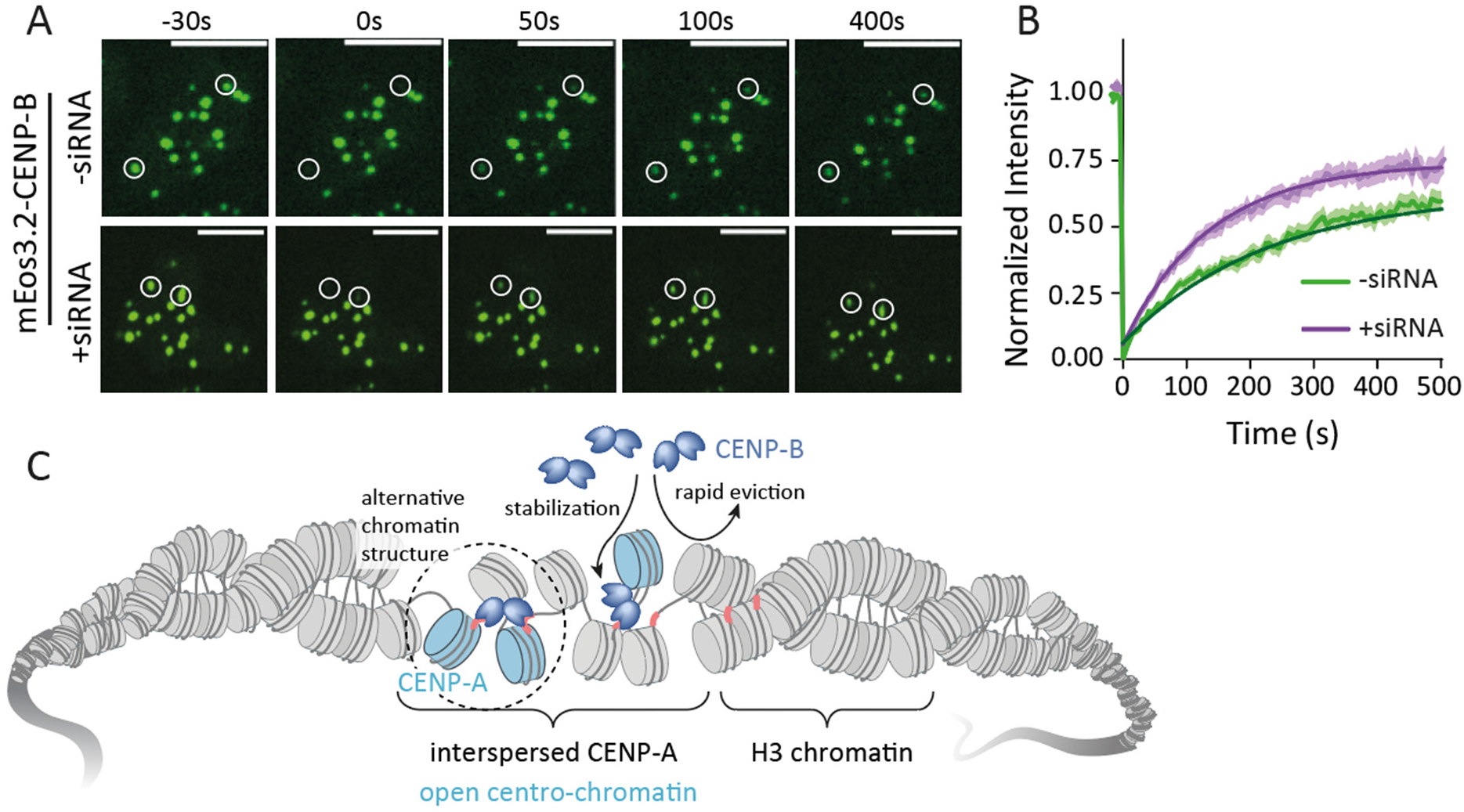
FRAP analysis of CENP-B in the presence and absence of CENP-A depletion in interphase cells. **A**) Representative FRAP analysis of mEos3.2-CENP-B in the presence (top, -siRNA) or absence (bottom, 48 hrs after siRNA treatment) of CENP-A. Bleached areas are indicated with white circles. T<0 frame is before bleaching and signal recovery is shown at indicated time points. **B**) Normalized recovery curves for quantitative FRAP measurements of CENP-B with or without CENP-A depletion. Thick lines indicate mean values and the colored areas indicate the standard error. At least 10 cells and 20 centromeres were used for analysis. **C**) Model of the collaboration between CENP-A and CENP-B to generate an open centro-chromatin state.

Together, our results show that CENP-B chromatin interaction dynamics depend on the presence of CENP-A, creating a permissive chromatin environment (**Figure 6C**). Moreover, stable CENP-B localization at centromeres is impaired in C-terminal truncations, indicating that protein regions outside of the DNA binding domain contribute to stable chromatin localization, and that protein dimerization is important for CENP-B function.

## Discussion

CENP-A-containing nucleosomes lie at the core of centromeric chromatin organization, as they form an interaction hub for the CCAN and kinetochore complexes^13,59,60^. Several centromere proteins, including CENP-C, CENP-N and CENP-B, have been shown to directly contact CENP-A nucleosomes, and thereby may act as anchors for the kinetochore on chromatin^8,14^. This potentially requires a distinct chromatin organization, enabling such complex multivalent interactions.

Here, we investigated the dynamic centromeric chromatin organization using single-molecule fluorescence approaches. Measuring FRET between neighboring nucleosomes, we observed that CENP-A renders chromatin structure open and highly dynamic. Next-neighboring stacking interactions between nucleosomes were reduced and highly transient, revealing large-scale fluctuations on the seconds timescale, compared to the more rapid, local and small-amplitude conformational fluctuations within H3 chromatin^39^. However, our FRET measurements also revealed that the linker DNA extending from CENP-A nucleosomes was ordered within a fiber context, in contrast to the greatly flexible DNA linkers observed in single nucleosomes^15,16^. Together, these results show that CENP-A containing chromatin can adopt a regular higher-order structure, but due to the flexibility of CENP-A nucleosomes, nucleosome stacking interactions are transient and fiber access is greatly increased. This effect is most likely further pronounced in native centromeres, as actual centromeric DNA such as alpha-satellite repeats in humans are less strongly nucleosome bound, compared to engineered 601 DNA. In particular, human centromeric DNA can form alternative hairpin structures, which most likely further distort chromatin structure^38^.

Centromere proteins that require direct access to CENP-A nucleosomes or centromeric DNA, such as CENP-B, compete against nucleosome-nucleosome contacts within a chromatin fiber, resulting in increased dissociation kinetics. CENP-A chromatin with its increased flexibility releases this inhibition, restoring unhindered DNA access to CENP-B. While direct interactions between CENP-B and CENP-A have been proposed^33^, we could not however detect direct stabilization of CENP-B binding via CENP-A-mediated interactions in a mononucleosome context. Conversely, CENP-B itself further opens chromatin structure upon binding and destabilizes nucleosome-stacking interactions. The two centromeric proteins thus collaborate to establish a more accessible centromeric chromatin state. Nevertheless, CENP-B binding is only hindered but not excluded from H3 chromatin. In agreement with these observations, in cells, CENP-B is found throughout the centromere and is not limited only to CENP-A nucleosomes^23,61^. Still, when depleting CENP-A from cellular chromatin, we observed an increase in the dynamics of CENP-B, indicating weaker chromatin binding. This is in agreement with earlier studies that showed a loss of CENP-B signal at centromeres in CENP-A-depleted cells^61^, underlining the fact that CENP-B retention at the centromere is reduced by CENP-A loss.

Thus, increased flexibility of CENP-A-containing chromatin is likely an important factor for CCAN recruitment by allowing biochemical access to both the CENP-A nucleosome as well as the linker DNA. This is important as recent cryo-EM studies revealed a structured CCAN complex, which may induce considerable conformational changes within chromatin when bound^21,62^. In contrast, the CCAN member CENP-N can induce the compaction of CENP-A-containing arrays^63^ by bridging and thereby stabilizing nucleosome interactions, and overcoming the inherent flexibility of CENP-A chromatin. It is, therefore, still an open question of how chromatin structure at the centromeres is coupled to its function. Interestingly, while CENP-N stabilizes interphase recruitment of key centromere proteins such as CENP-C ^8,64^, its levels are markedly reduced at the centromere during kinetochore assembly during mitosis, correlated with structural changes in centro-chromatin^65,66^. Together, our studies show how histone variants such as CENP-A can reorganize chromatin structure, and thus promote chromatin invasion by DNA binding proteins, including CENP-B. This sets the stage for large-scale organization of centromeric chromatin, including the formation of long-range loops stabilized by CENP-B dimerization. The elastic and dynamic chromatin state, established by CENP-A and CENP-B, then allows the binding of CCAN members, attachment of the kinetochore and buffering of the forces exerted upon chromosomes during cell division.

## Acknowledgments

We thank Karolin Luger and Keda Zhou for CENP-A tetramers, and initial discussions on the project, Daniele Fachinetti for the donation of the DLD-1 cell line, and Maxime Mivelaz and Ruud Hovius for comments on the manuscript. BF thanks the Swiss National Science Foundation (Grant no. 31003A_173169), the European Research Council (ERC Consolidator grant no. 724022) and EPFL for financial support.

