## Supplementary Material for "CENP-A and CENP-B collaborate to create an open centromeric chromatin state"

[1]École Polytechnique Fédérale de Lausanne (EPFL), SB ISIC LCBM, Station 6, CH-1015 Lausanne, Switzerland

### In-vitro biochemical experiments

#### Expression and purification of JF549-CENP-B

The ybbR-CENP-B-$His_6$, and ybbR-CENP-$B_{1-150}$-$His_6$ constructs were cloned into pET-15b and expressed in Rosetta-Gami 2 E. coli cells. Typically, 1L cultures of bacteria was grown to OD = 0.8 (600 nm) at 37° C before induction with 50 µg/mL IPTG and subsequently left overnight at 16° C with shaking. Bacteria were then harvested by centrifugation (3000 x g, 5 mins), resuspended in 10 mL/L culture of resuspension buffer (phosphate buffered saline (PBS, 137 mM NaCl, 2.7 mM KCl, 8 mM $Na_2HPO_4$, and 2 mM $KH_2PO_4$) supplemented with 500 mM NaCl, and protease inhibitors (Roche)), and lysed by sonication. Post-sonication cell debris was removed by centrifugation (100’000 x g, 45 mins) and the resulting supernatant was filtered through a 0.2 µm filter.

The cleared cell lysate was passed through a 5 mL Histrap column (GE, AKTA system) pre-equilibrated in resuspension buffer. The His-column was washed in wash buffer 1 (10 mM Tris-HCl (pH 7.5), 2 M NaCl, 1 mM DTT) to remove any DNA contamination and wash buffer 2 (10 mM Tris-HCl (pH 7.5), 500 mM NaCl, 1 mM DTT, and 20 mM Imidazole) to remove non-specifically bound proteins until a stable UV signal was seen on the FPLC system. The column was then eluted using a 10 CV (column volume) gradient elution from wash buffer 2 to elution buffer (10 mM Tris-HCl (pH 7.5), 500 mM NaCl, 1 mM DTT, and 500 mM Imidazole). Fractions containing CENP-B were identified by SDS-PAGE and the presence of CENP-B protein was confirmed by immune-blotting. Fractions were pooled and concentrated to ~250 µL via Amicon 50K MWCO centrifugal spin filters. The concentrated protein was then dialyzed into storage buffer (10 mM Tris-HCl (pH 7.5), 500 mM NaCl, 1 mM DTT, 10% glycerol) and the final concentration was determined by UV spectroscopy.

For fluorescent labelling, CENP-B was diluted to a final concentration of 5 µM in reaction buffer (50 mM HEPES (pH 7.5), 10 mM $MgCl_2$, 1 µM SFP synthase, 10 µM CoA-JF549) and incubate overnight at 4° C. The reaction mixture was centrifuged at 13’000 rpm, 5 min using a tabletop centrifuge (Eppendorf) to remove aggregates before performing size exclusion purification on a Superdex200 10/300 or Superdex75 10/300 column (GE) for CENP-B or CENP-$B_{1-150}$ respectively.

Fractions were analyzed using SDS PAGE. Clean samples were pooled, concentrated, and flash frozen for storage at -80° C. Labeling efficiency and concentration were measured by UV spectroscopy. The labeling efficiency with JF549 dye was approx. 50%, as judged from UV-VIS spectroscopy (**Figure S3**).

#### Expression and purification of recombinant histones

CENP-A tetramers were purified from E. Coli as described[1,2], or purchased from EpiCypher. Canonical histones were expressed and purified as described before[3]. Briefly, individual wild-type human histones were cloned into pET-15b plasmid vectors and expressed in BL21 DE3 plysS cells. Cells were grown in LB media containing 100 µg/mL ampicillin and 35 µg/mL chloramphenicol at 37° C until OD600 = 0.6. Expression was induced by IPTG addition to a final concentration of 0.5 mM. After 3 h expression, cells were harvested by centrifugation and resuspended in lysis buffer (20 mM Tris pH 7.5, 1 mM EDTA, 200 mM NaCl, 1 mM βMe, Roche protease inhibitor) and frozen. Cells were lysed by freeze-thawing and sonication. Inclusion bodies were harvested by centrifugation. The inclusion body pellet was washed once with 7.5 mL of lysis buffer containing 1% Triton and once without. Inclusion body pellets were resolubilized in resolubilization buffer (6 M GdmCl, 20 mM Tris pH 7.5, 1 mM EDTA, 1 mM β-mercaptoethanol (βMe)) and dialyzed into urea buffer (7 M urea, 10 mM Tris, 1 mM EDTA, 0.1 M NaCl, 5 mM 1 mM βMe, pH 7.5). Histones were purified by cation exchange chromatography using a HiTrap SP HP 5 mL column (GE Healthcare). Fractions were analyzed by SDS-PAGE and pooled, followed by dialysis into water and lyophilization. Final purification was performed by preparative RP-HPLC. Purified histones were lyophilized and stored at −20° C until used for octamer refolding.

**Dimer and octamer refolding**

In a typical octamer refolding reaction, 0.4 mg of each of the pure lyophilized human histones were dissolved in unfolding buffer (6 M GdmCl, 10 mM Tris, 5 mM DTT, pH 7.5) to an expected concentration of 2 mg/mL. The exact concentration was determined by UV spectroscopy, using the following extinction coefficients: $\varepsilon_{280nm,H2A} = 4470\ M^{-1}cm^{-1}$, $\varepsilon_{280nm,H2B} = 7450\ M^{-1}cm^{-1}$, $\varepsilon_{280nm,H3} = 4470\ M^{-1}cm^{-1}$, $\varepsilon_{280nm,H4} = 5960\ M^{-1}cm^{-1}$. For dimer refolding, equimolar amounts of H2A and H2B were mixed and unfolding buffer was added to a final histone concentration of 1 mg/mL. For octamer refolding, equimolar amounts of H3 and H4 were mixed along with 1.05 equivalents of H2A and H2B and unfolding buffer was added to a final histone concentration of 1 mg/mL. Dimers or octamers were then refolded by dialysis against refolding buffer (2 M NaCl, 10 mM Tris, 1 mM EDTA, 5 mM DTT, pH 7.5). The refolded dimers or octamers were subsequently purified by gel filtration on a Superdex S200 10/300GL column. Collected fractions were analyzed by SDS-PAGE, and octamer containing fractions were pooled and concentrated. Finally, glycerol was added to a final concentration of 50%, concentrations were determined by UV spectroscopy and octamer stocks were stored at -20° C.

**Oligonucleotide labeling**

Fluorescently labeled oligonucleotides were generated as described before[4]. Briefly, 5-10 nmol of single-stranded oligonucleotide, containing amino modifier C6 dT (purchased from Integrated DNA Technologies IDT), was diluted in 25 µl labeling buffer (0.1 M sodium tetraborate, pH 8.5). 5 µl of a 5 mM stock of succinimidyl-ester modified fluorophore (Alexa 647 or Cy3B) were added to the reaction mix and left shaking at 4° C overnight. For a table enumerating all labeled oligonucleotides see **Table S2**. The reaction progress was monitored by RP-HPLC using a gradient from solvent A (95% 0.1M triethylammonium acetate (TEAA) pH 7, 5% acetonitrile) to solvent B (70% 0.1M TEAA pH 7, 30% acetonitrile) on a 3 µm 4.6 x 150 mm InertSustain C18 column (GL sciences) over 20 min. More dye was added when required. For purification, the labeled DNA was precipitated via the addition 0.3M NaOAc pH 5.2 followed by 2.75 equivalents of cold ethanol (ethanol precipitation), followed by centrifugation at 20'000 x g at 4° C for 20 min. This was repeated twice to remove excess unconjugated dye. The DNA pellet was finally dissolved in 100 µl solvent A and purified by HPLC. The purified DNA was finally isolated by ethanol precipitation and dissolved in milliQ water to a concentration of ~10 µM.

**Preparation of mononucleosomes**

Labelled nucleosome DNA was prepared by PCR (fragment 1x601 and 1x601BB. Sequences in **Table S2**). Nucleosomes (H3 containing 1x601, 1x601BB, and CENP-A containing 1x601, 1x601BB) were prepared following ref. [5]. Typically, 1-5 µg of labeled and biotinylated DNA (1x601, 1x601BB) was combined with purified refolded H3 containing histone octamers or a mixture of 2 eq. H2A/H2B dimers and 0.9 CENP-A/H4 tetramers at experimentally determined ratios (1:1 to 1:3, DNA to histone octamer ratio) in 20 µl TE (10 mM Tris-HCl pH 7.5, 1 mM EDTA) supplemented with 2 M KCl. In a typical dialysis, the mixture was added to a micro-dialysis unit (Thermo Scientific, Slide-A-Lyzer – 10'000 MWCO), then dialyzed into TE buffer (10 mM Tris pH 7.5, 0.1 mM EDTA pH 8.0) with a linear gradient from 2 M to 10 mM KCl for 16-18 h, and finally kept in TEK10 buffer (10 mM Tris pH 7.5, 0.1 mM EDTA pH 8.0, 10 mM KCl) for another 1 h. Samples were then centrifuged at 20'000 x g for 10 min at 4° C and the supernatant was kept on ice. To determine the quality of MN assemblies, 5% polyacrylamide gels were analyzed using native PAGE in 0.25 x TB at 90 V on ice for 90 min (**Figures S3A,B**).

**Electrophoretic mobility shift assays (EMSA)**

EMSAs to determine CENP-B binding to DNA or mononucleosomes were performed in single-molecule imaging buffer (10 mM Tris pH 7.5, 130 mM KCl, 3.2% w/v glucose), with 20 µL total volume. Typically, 200 nmol stocks of DNA and 3 µM stocks of CENP-B were prepared and serially diluted to desired concentrations. Reactions were mixed by pipetting and incubated for 30 min at room temperature. Sucrose was added to a final concentration of 8% and reactions were loaded onto 5% polyacrylamide gels run in 0.25 x TBE at 90 V for 90 min. Images were taken using ChemiDoc MP (Biorad) (**Figures S5**).

**Plug and Play synthesis of labelled 12x601 DNA**

Singly-labeled and biotinylated 12x601 DNA was produced as shown in **Figure S1A**. In short, recombinant 12x601 arrays were generated containing sites for nicking endonucleases Nb.BssSI and Nt.BstNBI (NEB) (sequences in **Table S1**). Typically, 100 µg of 12x601 containing plasmids was digested with EcoRV, HindIII-HF, and Calf alkaline phosphatase in Cutsmart buffer (NEB) to release the 12x601 DNA. The DNA fragments were then purified by PEG precipitation, followed by PEG removal using Qiaquick PCR purification spin columns (Qiagen) to generate ~50 µg of array DNA. The arrays were subsequently treated with Nb.BssSI and Nt.BstNBI (NEB) at 50° C for 2 hrs to generate nicked sites. 10-fold excess of complimentary oligonucleotides (**Table S2**) were added to the nicked arrays and annealed by heating to 90° C and cooling by 1° C/min until 25° C. Nicked sites on annealed arrays were sealed by T4 DNA ligase treatment (NEB) for 1hr in Cutsmart buffer supplemented with ATP. Biotin-containing anchors were ligated to the HindIII site. Labelling was checked by agarose gel electrophoresis (**Figure S1A**) and labelled array DNA was purified by PEG precipitation and PCR purification spin columns. Final concentration was determined by UV spectroscopy. Typical yield for 100 µg starting plasmid was ~20 µg of labelled arrays.

**Reconstitution of 12x601 chromatin fibers**

Labelled array DNA was prepared by plug and play synthesis as described above (Sequences in **Table S1-3**). Chromatin arrays (H3 containing chromatin on 12x601 DA1, DA1BB, DA3BB DNA (BB: B-B-Box), and CENP-A containing chromatin on 12x601 DA1BB, 12x601DA3BB DNA) were prepared following ref. [5]. Typically, 100-200 pM of labeled and biotinylated 12x601 DNA was combined with purified refolded H3 containing histone octamers or a mixture of 2 eq. H2A/H2B dimers and 0.9 CENP-A/H4 tetramers at experimentally determined ratios (1:1 to 1:3, DNA to histone octamer ratio) in 30 µl TE (10 mM Tris-HCl pH 7.5, 1 mM EDTA) supplemented with 2 M KCl. H3 containing arrays were also supplemented with 0.5-1 eq. MMTV DNA to prevent octamer oversaturation. In a typical dialysis, the mixture was added to a micro-dialysis unit (Thermo Scientific, Slide-A-Lyzer – 10'000 MWCO), then dialyzed into TE buffer (10 mM Tris pH 7.5, 0.1 mM EDTA pH 8.0) with a gradient from 2 M to 10 mM KCl for 16-18 h, and finally equilibrated in TEK10 buffer (10 mM Tris pH 7.5, 0.1 mM EDTA pH 8.0, 10 mM KCl) for another 1 h. Samples were then centrifuged at 20'000 x g for 10 min at 4° C and the supernatant was kept on ice. To determine the quality of array assemblies, they were analyzed using native PAGE on 0.6% Agarose 0.25 x TB gels, run at 90 V on ice for 90 min (**Figures S1B-C**). Chromatin assembly quality was further verified by ScaI digestion of 12x assemblies. Only samples showing full saturation and minimal free 601 DNA were used for further experiments.

**Preparation of microfluidic chambers for sm-FRET/TIRF experiments**

Cleaning, silanization and PEGylation of coverslips and glass slides was done as described previously[3]. Briefly, coverslips (24 × 40 mm, 1.5 mm thickness) and glass slides (76 × 26 mm with 2 rows of 4 holes drilled) were sonicated for 20 min in 10% Alconox, rinsed with milliQ water and the procedure was repeated sequentially with acetone and ethanol. Both coverslips and glass slides were then placed in piranha etching solution (25% v/v 30% $H_2O_2$ and 75% v/v $H_2SO_4$) for a minimum of 2 h. After thorough washing with milliQ $H_2O$, coverslips and slides were sonicated in acetone for 10 min, then incubated with 2% v/v aminopropyltriethylsilane (APTES) in acetone for 15 min, and dried. Flow-chambers were assembled from one glass slide and one coverslip separated by machine-cut double-sided 0.12 mm tape (Grace Bio-labs) with channels between each opposing hole in the glass slide. Pipette tips were fitted in each of the 2 × 8 holes on each side of the silanized glass flow chambers as inlet reservoir and outlet sources and glued in place with epoxy glue. The glue was allowed to solidify for 30 - 40 min. Subsequently, 350 µL of 0.1 M tetraborate buffer at pH 8.5 was used to dissolve ~1 mg of biotin-mPEG(5000 kDa)-SVA (Layson Bio), and 175 µL from this was transferred to 20 mg mPEG (5000kDa)-SVA. This was centrifuged and mixed to homogeneity with a pipette before 10-20 µL aliquots were loaded into each of the eight channels in the flow chamber. The PEGylation reaction was allowed to continue for the next 2½-4 h after which the solution was washed out with degassed ultra-pure water (Romil).

**Single-molecule TIRF (sm-TIRF) co-localization microscopy measurements**

Measurements were done according to ref.[3]. Objective-type smTIRF was performed using a fully automated Nikon Ti-E inverted fluorescence microscope, equipped with an ANDOR iXon EMCCD camera and a TIRF illuminator arm, controlled by NIS-elements and equipped with a CFI Apo TIRF 100 x oil immersion objective (NA 1.49), resulting in a pixel size corresponding to 160 x 160 nm. Laser excitation was realized using a Coherent OBIS 640LX laser (640 nm, 40 mW) and Coherent OBIS 532LS laser (532 nm, 50 mW) on a custom-made laser bench. Wavelength selection and power modulation was done using an acousto-optical tunable filter (AOTF) controlled by NIS-elements. Typical laser intensities in the objective used for measurements were 0.8 mW for both 532 nm and 640 nm laser lines. For all smTIRF experiments, flow channels were washed with 100 µL degassed ultrapure water (Romil), followed by 100 µL 1 x T50 (10 mM Tris pH 8, 50 mM NaCl). Channel quality was checked by observing background fluorescence using both 532 nm and 640 nm excitation. 50 µL of 0.2 mg/mL neutravidin was then injected and incubated for 5 min, followed by washes using 200 µL of 1 x T50 buffer. 50 pM of Alexa647 labeled DNA/mononucleosomes/12-mer chromatin assemblies were then flowed in for immobilization in T50 with 2 mg/mL bovine serum albumin (BSA, Carlroth). A 25 × 50 µm imaging area was monitored using 640 nm excitation to check for sufficient coverage. 200 µL 1 x T50 was used to wash out unbound Alexa647 labeled DNA/mononuceosomes/12-mer chromatin assemblies. 5-10 nM JF-549 labeled CENP-

B or CENP-$B_{1-150}$ (see table below for details) was flowed in using imaging buffer (50 mM Tris pH 7.5, 40 mM KCl, 110 mM NaCl 10% v/v glycerol, 0.005% v/v Tween 20, 2 mM Trolox, 2 mM nitrobenzyl alcohol (NBA), 2 mM cyclooctatetraene (COT), 3.2% w/v glucose, 1x glucose oxidase/catalase oxygen scavenging system and 2 mg/mL BSA). Images were recorded using the following parameters where $t_{on}$ denotes the camera integration time, and $t_{off}$ indicates interspersed time intervals of camera inactivity:

| | Camera $t_{on}$ (msec) | Camera $t_{off}$ (msec) | Orange channel # frames | Far-red channel: # frames | n repeat |
|---|---|---|---|---|---|
| DNA | 100 | 0.3 | 99 | 1 | 40 |
| Mononucleosome | 100 | 0.3 | 99 | 1 | 40 |
| 12-mer chromatin fiber | 100 | 0.3 | 99 | 1 | 40 |

**Colocalization data analysis**

Single-molecule trace extraction and trace analysis were done according to ref. [3,6] with some adjustments. Acquired movies were background-corrected in ImageJ using a rolling ball algorithm. Trace extraction and analysis was performed in custom-written MATLAB software: DNA/nucleosome or chromatin positions were detected via a local maxima approach. Sequential images were aligned using the far-red channel to compensate for stage drift. Fluorescence intensities (in the orange channel) were extracted from the stack within a 2 pixel radius of the identified DNA peaks. Every detected spot in the orange channel was fitted with a 2D-Gaussian function to determine co-localization with immobilized DNA/chromatin. Peaks exceeding an experimentally determined PSF width for a single JF-549 molecule were excluded from further analysis. For noise reduction, extracted fluorescence traces were filtered using a forward-backward non-linear filter[7].

Individual binding events were detected using a thresholding algorithm. Overlapping multiple binding events were excluded from the analysis. For each movie, cumulative histograms were constructed from detected bright times ($t_{bright}$) corresponding to bound CENP-B molecules. The cumulative histograms from traces corresponding to individual DNA / mononucleosome / chromatin fibers were fitted with di-exponential functions:

$$y = \sum_{i=1}^{2} A_i \exp(-t / \tau_{off,i})$$

yielding non-specific residence times $\tau_{off,0}$ or the specific residence times $\tau_{off,1}$ and $\tau_{off,2}$ . Cumulative histograms constructed from dark times ($t_{dark}$), in between binding events, were fitted with mono-exponential functions:

$$y = A \exp(-t \cdot k_{on,app})$$

to obtain apparent on-rate constants.

**Single-molecule FRET (smFRET) measurements**

Flow cell preparation and chromatin loading was performed as described in ref.[4,8] and the preceding paragraphs. Experiments were performed in FRET imaging buffer (40 mM KCl, 50 mM Tris pH 7.5, 2 mM Trolox, 2 mM nitrobenzyl alcohol (NBA), 2 mM cyclooctatetraene (COT), 10% glycerol and 3.2% glucose) supplemented with GODCAT (100 x stock solution: 165 U/mL glucose oxidase, 2170 U/mL catalase). Experiments on the effect of CENP-B on chromatin compaction were supplemented with 2 mg/ml BSA to prevent non-specific interactions. smFRET data acquisition was carried out with a micro-mirror TIRF system (MadCityLabs) using Coherent Obis Laser lines at 405 nm, 488 nm, 532 nm and 640 nm, a 60x NA 1.49 Nikon CFI Apochromat TIRF objective (Nikon) as well as an iXon Ultra EMCCD camera (Andor), operated by custom-made Labview (National Instruments) software. For general smFRET imaging, a programmed sequence was employed to switch the field of view to a new area followed by adjusting the focus. Subsequently 2000 frames at 100 ms integration time were recorded under alternating excitation (ALEX) with using a 532 nm and 640 nm laser line. Each experiment ended with a strong laser pulse in the donor and acceptor channel to bleach the dyes for better background determination. Moreover, each experiment was repeated several times (see **Tables S4** and **S6** for number of repeats), using at least two independently produced chromatin preparations on two different days.

**FRET data analysis**

Similar to colocalization experiments, acquired movies were background-corrected in ImageJ, whereas trace extraction and analysis was performed in custom-written MATLAB software. Single-molecule fluorescence peaks were automatically detected in the initial acceptor image prior to donor excitation and the same peaks were selected in the donor channel. The donor and the acceptor images were aligned using a transformation matrix generated from 4-6 peaks appearing in both the donor and the acceptor channels. Peaks tightly clustered, close to the edges or above a set intensity threshold in either the donor or the acceptor channels were removed from analysis. All remaining traces were extracted for further analysis. Using the ALEX data, traces that showed less or more than exactly one donor or acceptor dye (due to clustering of molecules or incomplete labeling) were excluded from the analysis.

From traces of donor- ($F_D$) and acceptor ($F_A$) fluorescence emission intensity, FRET efficiency ($E_{FRET}$) traces are calculated as follows:

$$E_{FRET} = \frac{F_A - \beta F_D}{F_A - \beta F_D + \gamma F_D} \text{ where } \beta = \frac{F_{A,bleach}}{F_{D,bleach}} \text{ and } \gamma = \frac{\Delta F_{A,bleach}}{\Delta F_{D,bleach}}$$

where β denotes the bleed-through of acceptor fluorescence into the donor channel, and γ is the sensitivity ratio of the detection system for donor and acceptor fluorescence. The values of $\beta = 0.071$ and $\gamma = 0.463$ were experimentally determined for the dye pair Cy3B/Alexa647 in our experimental setup. From fluorescence time-traces, $E_{FRET}$ histograms were constructed, using a bin size of 0.02. $E_{FRET}$ histograms of each trace of length > 5 s were normalized to total counts. Final histograms for each independent measurement were fitted using 2 or 3 Gaussian functions $\sum_i A_i e^{-\left(\frac{x-c_i}{\sigma_i}\right)^2}$. For fit values, see **Tables S4** and **S6**.

Traces were selected based on the following criteria: 1) Initial total fluorescence of the donor and the γ-corrected acceptor of > 2000 counts over baseline (at 900 EM gain). 2) At least 5 s prior to bleaching of acceptor or donor. 3) Single bleaching event for donor or acceptor. 3. a) if acceptor bleaches first; leads to anticorrelated increase in donor to same total fluorescence level as prior to bleaching. 3.b) if donor bleaches first, the acceptor dye must still be fluorescent when directly excited. Traces that did not match these criteria were rejected from the analysis. Traces were finally sorted into 'static' and 'dynamic' dependent on the existence of anticorrelated intensity fluctuations in donor- and acceptor channels.

### Cell experiments

#### Cell line construction and culture conditions

The mEos3.2-CENP-B/-CENP-$B_{1\text{-}500}$/-CENP-$B_{1\text{-}150}$ constructs were generated by subcloning into pcDNA5/FRT/TO vector (Thermo-Fischer) by Gibson cloning. In short, the mEos3.2 and CENP-B coding regions were created using primers with complementary 3' and 5' overhangs, respectively to each other and 5' and 3' respectively with pcDNA/FRT/TO. These were mixed with PCR linearized pcDNA5/FRT/TO and infusion mix (Takara clonetech) to generate the final constructs.

Cell lines were generated using lipofectamine 2000 (Invitrogen) based transfection as per the manufacturer's guidelines. In short, to generate cell lines, $1 \times 10^5$ DLD-1 FLP-In cells were grown in 24 well plates until 70-90% confluent. Subsequently 0.7-2 µg of CENP-B-pcDNA5/FRT/TO was mixed with 2.5-8 µg of pOG44 (Thermo-Fischer) in 500ul of serum-free medium and 10ul lipofectamine 2000. The mixture was allowed to sit for 20 mins to form DNA-lipid vesicles. The mixture was added to the cells for 6 hrs before the medium was changed to 10% FBS-DMEM for 16 hrs. After 24 hrs 500 µg/mL hygromycin was added to perform selection. Individual colonies were picked after 1 month and grown in hygromycin. Expression of constructs was checked by immune-blotting and immunofluorescence.

All cell lines were grown in DMEM supplemented with 10% fetal bovine serum (FBS) and 100 U/mL Penicillin-Streptomycin and tested for mycoplasma (using EZ-PCR, Biological industries).

#### Immunofluorescence and immune-blotting

Human DLD-1 mEos3.2-CENP-B/-CENP-$B_{1\text{-}500}$/-CENP-$B_{1\text{-}150}$ cells were grown on glass bottom 12 well plates (MatTek) until 70% confluent and fixed in 3% paraformaldehyde for 10 mins at room temperature. Staining of cells was performed using rabbit anti-CENP-C antibodies (Abcam ab193666) as the primary antibody and Donkey anti-rabbit IgG AF647 (Abcam ab150075). Images were collected on an inverted stage Nikon microscope with a 100 x oil immersion objective. All images were analyzed in ImageJ.

Western blotting was conducted using either a Trans-blot SD semi dry transfer cell (Bio-Rad) or an iBlot 2 transfer system (Invitrogen). Western blots were performed against CENP-A (2186S, Cell Signaling), CENP-B (ab25734, Abcam), and alpha-tubulin (T9026, Sigma-Aldrich).

**FRAP analysis of CENP-B DLD-1 cell lines**

Human DLD-1 mEos3.2-CENP-B/-CENP-$B_{1-500}$/-CENP-$B_{1-150}$ cells were grown until 70% confluent and then trypsinized for siRNA treatment. siRNA transfections were done with a mixture of 3 DsiRNAs (IDT, sequences in **Table S7**) or negative control DsiRNA (IDT) using the neon electroporation system (Invitrogen). In short, $2x10^5$ cells were mixed with 100-200 nM anti CENP-A or negative control DsiRNA and electroporated as per manufacturer recommendations. Cells were grown on glass bottom 12 well plates (MatTek) in DMEM supplemented with 10% fetal bovine serum (FBS) for 24 hrs before washing with PBS to remove dead cells. Cells were grown for a subsequent 24 hrs in DMEM supplemented with 10% fetal bovine serum (FBS) and doxycycline.

Microscopy was performed 48 hrs after transfection on a Nikon inverted stage spinning disk confocal microscope (Yokugawa CSU W1) with temperature and $CO_2$ stabilization at 37° C and 5% $CO_2$. Imaging was done using a Photometrics Prime 95B sCMOS and using a 60x oil immersion objective. A solid-state laser at 488 nm and 10-20 mW power was used for excitation. 1200 x 1200 pixels images at 16-bit color depth were acquired, using an imaging rate of 5 s per frame. Generally, 10 image frames were acquired before turning on the bleaching laser. Then, up to 16 circular spots of 5 pixels in diameter were used to bleach up to 2 centromeres in 8 cells. Background correction was performed using average intensity of cytosolic regions, followed by a photobleaching correction using the intensity unbleached centromeres (**Figure S9A**). Intensity graphs were normalized between the average pre-bleach intensity (1.0) and the first image after the bleach pulse (0.0). Results were averaged over ~30 FRAP curves for at least 10 cells. All images were taken using Nikon elements AR (Nikon) and analyzed by custom MATLAB code (available on request). All graphs were generated using GraphPad Prism 9.3.1 (GraphPad Software).

### Supplementary Tables

#### Table S1

| Fragment | Sequence |
|---|---|
| 1x601 | -100 -95 -90 -85 -80 -75 -70 -65 -60 -55 -50 -45 -40 -35 -30 -25<br>GATCGGTCTCATAGC**CTGGAGAATCCCGGTGCCGAGGCCGCTCAATTGGTCGTAGACAGCTCTA**<br>-20 -15 -10 -5 0 5 10 15 20 25 30 35 40 45 50 55<br>**GCACCGCTTAAACGCACGTACGCGCTGTCCCCCGCGTTTTAACCGCCAAGGGGATTACTCCCTAGTCTCCAGGCACGTGT**<br>60 65 70 75 80 85 90 95 100 105<br>**CAGATATATACATCCTGTCACACTGTGGATC** |
| 1x601 BB | -100 -95 -90 -85 -80 -75 -70 -65 -60 -55 -50 -45 -40 -35 -30 -25<br>GATCGGTCTCATAGCCTTCGTTGGAAACGGGA**GAATCCCGGTGCCGAGGCCGCTCAATTGGTCGTAGACAGCTCTA**<br>-20 -15 -10 -5 0 5 10 15 20 25 30 35 40 45 50 55<br>**GCACCGCTTAAACGCACGTACGCGCTGTCCCCCGCGTTTTAACCGCCAAGGGGATTACTCCCTAGTCTCCAGGCACGTGT**<br>60 65 70 75 80 85 90 95 100 105<br>**CAGATATATACATCCTGT**CACACTGTGGATC |
| P1 DA1 BB | -100 -95 -90 -85 -80 -75 -70 -65 -60 -55 -50 -45 -40 -35 -30 -25<br>CACTTGGTGGCGGCCGCC**CTGGAGAATCCCGGTGCCGAGGCCGCTCAATTGGTCGTAGACAGCTCTA**<br>-20 -15 -10 -5 0 5 10 15 20 25 30 35 40 45 50 55<br>**GCACCGCTTAAACGCACGTACGCGCTGTCCCCCGCGTTTTAACCGCCAAGGGGATTACTCCCTAGTCTCCAGGCACGTGT**<br>60 65 70 75 80 85 90 95 100 105<br>**CAGATATATACAAGATCT**AGTACTTGGTCTCATAGC |
| P1 DA3 BB | -100 -95 -90 -85 -80 -75 -70 -65 -60 -55 -50 -45 -40 -35 -30 -25<br>CACTTGGTGGCGGCCGCC**CTGGAGAATCCCGGTGCCGAGGCCGCTCAATTGGTCGTAGACAGCTCTA**<br>-20 -15 -10 -5 0 5 10 15 20 25 30 35 40 45 50 55<br>**GCACCGCTTAAACGCACGTACGCGCTGTCCCCCGCGTTTTAACCGCCAAGGGGATTACTCCCTAGTCTCCAGGCACGTGT**<br>60 65 70 75 80 85 90 95 100 105<br>**CAGATATATACAAGATCT**TCTCTAGATCCATGGAGTACTTGGTCTCATAGC |
| P2 DA1 BB | -100 -95 -90 -85 -80 -75 -70 -65 -60 -55 -50 -45 -40 -35 -30 -25<br>CTTCGTTGGAAACGGGA**GAATCCCGGTGCCGAGGCCGCTCAATTGGTCGTAGACAGCTCTA**<br>-20 -15 -10 -5 0 5 10 15 20 25 30 35 40 45 50 55<br>**GCACCGCTTAAACGCACGTACGCGCTGTCCCCCGCGTTTTAACCGCCACGAGGATTACTCCCTAGTCTCCAGGCACGAGC**<br>60 65 70 75 80 85 90 95 100 105<br>**CAGATATATACATCCTGTCACACTGTGCCAA** |
| P2 DA3 BB | -100 -95 -90 -85 -80 -75 -70 -65 -60 -55 -50 -45 -40 -35 -30 -25<br>C**TGGAGAATCCCGGTGCCGAGGCCGCTCAATTGGTCGTAGACAGCTCTA**<br>-20 -15 -10 -5 0 5 10 15 20 25 30 35 40 45 50 55<br>**GCACCGCTTAAACGCACGTACGCGCTGTCCCCCGCGTTTTAACCGCCAAGGGGATTACTCCCTAGTCTCCAGGCTCGTGT**<br>60 65 70 75 80 85 90 95 100 105<br>**CAGATATATACATCCTGT**CACACTGTGCCAAGTACT |
| P3 DA1 BB | -100 -95 -90 -85 -80 -75 -70 -65 -60 -55 -50 -45 -40 -35 -30 -25<br>GTACTTACGCGGCCGCCC**TGGAGAATCCCGGTGCCGAGGCCGCTCAATTGGTCGTAGACAGCTCTA**<br>-20 -15 -10 -5 0 5 10 15 20 25 30 35 40 45 50 55<br>**GCACCGCTTAAACGCACGTACGCGCTGTCCCCCGCGTTTTAACCGCCAAGGGGATTACTCCCTAGTCTCCAGGCACGTGT**<br>60 65 70 75 80 85 90 95 100 105<br>**CAGATACTGCAGAGATCT**AGTACTTGGTCTCAAACC |
| P3 DA3 BB | -100 -95 -90 -85 -80 -75 -70 -65 -60 -55 -50 -45 -40 -35 -30 -25<br>CTTCGTTGGAAACGGGA**GAATCTCGTGGCCGAGGCCGCTCAATTGGTCGTAGACAGCTCTA**<br>-20 -15 -10 -5 0 5 10 15 20 25 30 35 40 45 50 55<br>**GCACCGCTTAAACGCACGTACGCGCTGTCCCCCGCGTTTTAACCGCCAAGGGGATTACTCCCTAGTCTCCAGGCACGTGT**<br>60 65 70 75 80 85 90 95 100 105<br>**CAGATACTGCAGAGATCT**CACGAGCCATGGAGTACTTGGTCTCAAACC |
| P4 DA1 BB | -100 -95 -90 -85 -80 -75 -70 -65 -60 -55 -50 -45 -40 -35 -30 -25<br>CTTCGTTGGAAACGGGA**GAATCCCGGTGCCGAGTCCGCTCAATTGGTCGTAGAGTCCTCTA**<br>-20 -15 -10 -5 0 5 10 15 20 25 30 35 40 45 50 55<br>**GCACCGCTTAAACGCACGTACGCGCTGTCCCCCGCGTTTTAACCGCCAAGGGGATTACTCCCTAGTCTCCAGGCACGTGT**<br>60 65 70 75 80 85 90 95 100 105<br>**CAGATATATACATCC**TGTCACGTCGTGCCAA |

| | |
|---|---|
| P4 DA3 BB | -100 -95 -90 -85 -80 -75 -70 -65 -60 -55 -50 -45 -40 -35 -30 -25<br>**CACGAGAATCCCGGTGCCGAGGCCGCTCAATTGGTCGTAGACAGCTCTA**<br>-20 -15 -10 -5 0 5 10 15 20 25 30 35 40 45 50 55<br>**GCACCGCTTAAACGCACGTACGCGCTGTCCCCCGCGTTTTAACCGCCAAGGGGATTACTCCCTAGTCTCCAGGCACGTGT**<br>60 65 70 75 80 85 90 95 100 105<br>**CAGATATATACATCC**TGTCACGTCGTGCCAA |
| P5 | -100 -95 -90 -85 -80 -75 -70 -65 -60 -55 -50 -45 -40 -35 -30 -25<br>GTACTTACGCGGCCGCC**CTGGAGAATCCCGGTGCCGAGGCCGCTCAATTGGTCGTAGACAGCTCTA**<br>-20 -15 -10 -5 0 5 10 15 20 25 30 35 40 45 50 55<br>**GCACCGCTTAAACGCACGTACGCGCTGTCCCCCGCGTTTTAACCGCCAAGGGGATTACTCCCTAGTCTCCAGGCACGTG**<br>60 65 70 75 80 85 90 95 100 105<br>**TCAGATACTGCAGAGATCT**CTAGATCCGGTCTCACTAA |

**Table S1 | Sequences of 1 x 601 pieces for recombinant and PCR-generated DNA pieces.** 601 sequences indicated in bold. The labeled base pairs are indicated in red. The numbering is given as number of base-pairs relative to the dyad in the 601 sequence. B-Box (CENP-B binding) sites are indicated in blue. From these sequences, chromatin fiber DNA is assembled as described in **Table S3**.

### Table S2

**Table S4 | Sequences of all labeled oligonucleotides**

| Description | Dye | Sequence |
|---|---|---|
| P2_anchor_pos39_rev | Alexa 647 | 5'-biotin-GATCCACAGTGTGACAGGATGTATATATCTGACACGTGCC TGGAGAC/iAmMC6T/AGGGAG-3' |
| P5_PNP_anchor_antisense_dye | Alexa 647 | 5'-ph-AGCTTAGTCTGC/iAmMC6T/CAGTACTCGTCGCTAGATCCATG GTCCGATTACGCGG-3' |
| P5_PNP_anchor_antisense | - | 5'-ph-AGCTTAGTCTGCTCAGTACTCGTCGCTAGATCCATGGTCCGATT ACGCGG-3' |
| P5_PNP_anchor_sens | - | 5'-biotin-CCGCGTAATCGGACCATGGATCTAGCGACGAGTACTGAGC AGACT-3' |
| P3_DA3BB_pos82_PNP_insert | Alexa 647 | 5'-ph-TCGTGGGTTTGAGACCAAGTACTCCA/iAmMC6T/GGC-3' |
| P2_DA3BB_pos88_PNP_insert | Cy3B | 5'-ph-TCGTGTCAGATATATACATCCTGTCACACTGTGCCAAG/iAmMC6T/ ACTCTTCGT TGGAAACGGAAGAATC-3' |
| P2_DA1_pos39_PNP_insert | Cy3B | 5'-ph-TCGTGCCTGGAGAC/iAmMC6T/AGGGAGTAATCC-3' |
| P4_DA1_pos-39_PNP_insert | Alexa 647 | 5'-Ph-CAATTGG/iAmMC6T/CGTAGAGTCCTCT-3' |

**Table S2 | Sequences of all labeled oligonucleotides.** Ph: 5' phosphorylation. iAmMC6T: internal amino-modified C6 dT linker, used for dye attachment.

### Table S3

| Experiment | Name | Backbone | Dye | Modification |
|---|---|---|---|---|
| CENP-B binding | 1x601 | P3_DA1BB | Alexa647 (39) | 3' biotin |
| | 1x601 BB | P3_DA3BB | Alexa647 (39) | 3' biotin |
| CENP-B chromatin binding | DA3BB | P1,P2;P3DA3BB;P4,P5 | P5_anchor: Alexa647 | 3' biotin |
| Chromatin FRET | CH DA1BB FRET | P1 ; P2DA1BB ; P3 ; P4DA1BB ; P5 | P2: Cy3B (39), P4: Alexa647 (-39) | 3' biotin |
| | CH DA3BB FRET | P1 ; P2 ; P3DA3BB ; P4 ; P5 | P2: Cy3B (88), P3: Alexa647 (82) | 3' biotin |

**Table S3 | Overview of all chromatin DNA with different combinations of labels.** The DNA is composed of sequence elements in the order as described under 'Backbone', using sequences from **Table S1**. For chromatin DNA, using the plug-and-play strategy (**Figure S1**), indicated labeled sequences (see **Table S2**) are swapped in and ligated.

### Table S4

| | | CENP-A DA1BB FRET | | CENP-A DA3BB FRET | | H3 DA1BB FRET | | H3 DA3BB FRET | |
|---|---|---|---|---|---|---|---|---|---|
| | | 40 mM KCl | 4 mM Mg2+ | 40 mM KCl | 4 mM Mg2+ | 40 mM KCl | 4 mM Mg2+ | 40 mM KCl | 4 mM Mg2+ |
| **LF** | A1 | 5.51 ± 0.8 | 4.9 ± 0.8 | 6.81 ± 0.42 | 3.72 ± 0.78 | 2.24 ± 1.18 | 2.52 ± 0.44 | 2.27 ± 0.4 | x |
| | c1 | 0.036 ± 0.022 | 0.056 ± 0.03 | 0.07 ± 0.02 | 0.074 ± 0.006 | 0.047 ± 0.035 | 0.05 ± 0.02 | 0.10 ± 0.03 | x |
| | s1 | 0.1 ± 0.03 | 0.1 ± 0.02 | 0.05 ± 0.004 | 0.06 ± 0.008 | 0.06 ± 0.03 | 0.12 ± 0.03 | 0.084 ± 0.022 | x |
| | % area | 77.74 ± 13.8 | 61.76 ± 6.46 | 39.78 ± 6.62 | 29.4 ± 9.68 | 19.67 ± 12.72 | 38.37 ± 2.18 | 16.03 ± 13.93 | x |
| **HF** | A2 | 1.28 ± 0.73 | 2.13 ± 0.3 | 3.43 ± 0.36 | 4.47 ± 0.67 | 6.57 ± 1.04 | 3.96 ± 0.59 | 7.04 ± 1.04 | 6.81 ± 0.7 |
| | c2 | 0.19 ± 0.05 | 0.44 ± 0.05 | 0.18 ± 0.02 | 0.32 ± 0.002 | 0.26 ± 0.036 | 0.57 ± 0.01 | 0.36 ± 0.035 | 0.47 ± 0.02 |
| | s2 | 0.12 ± 0.05 | 0.14 ± 0.03 | 0.13 ± 0.002 | 0.012 ± 0.004 | 0.09 ± 0.015 | 0.125 ± 0.016 | 0.09 ± 0.026 | 0.10 ± 0.016 |
| | % area | 22.25 ± 13.8 | 38.2 ± 6.46 | 60.21 ± 6.62 | 70.6 ± 9.68 | 80.32 ± 12.72 | 61.6 ± 2.18 | 83.96 ± 13.93 | 100 |
| **N total traces** | | 404 | 483 | 366 | 335 | 404 | 530 | 417 | 418 |
| **N dynamic traces** | | x | 129 | x | 53 | x | 32 | x | 15 |
| **% dynamic traces** | | x | 25.24 ± 4.72 | x | 15.6 ± 0.64 | x | 6.0 ± 2.3 | x | 2.9 ± 2.3 |
| **N of repeats** | | 4 | 4 | 2 | 2 | 4 | 3 | 3 | 3 |

**Table S4 | Summary of Gaussian fits and percentage of dynamic traces from chromatin compaction experiments.** Errors are fitting errors. N dynamic traces are determined by the existence of anticorrelated intensity fluctuations in donor- and acceptor channels.

### Table S5

| | dissociation kinetics | | | | binding kinetics | | |
|---|---|---|---|---|---|---|---|
| | dwell time (s) | | Amplitude (%) | | rate constants (x $10^7$ M*s-1) | | |
| | $\tau_{off,0}$ | $\tau_{off,1}$ | $A_0$ | $A_1$ | $k_{on}$ | $k_{on, per DNA repeat}$ | n exp. |
| **DNA** | | | | | | | |
| 1x 601 | 1.68 ± 0.5 | 20.89 ± 5.8 | 65 ± 8 | 34 ± 8 | 8.1 ± 4.6 | 8.1 ± 4.7 | 4 |
| 1x601 BB | 6.94 ± 1.42 | 61.76 ± 11.8 | 49 ± 9.5 | 51 ± 9.5 | 9.9 ± 2.5 | 9.9 ± 2.5 | 4 |
| **Nucleosomes** | | | | | | | |
| H3 1x601 MN | 3.4 ± 0.8 | 15.2 ± 1 | 51 ± 3.5 | 49 ± 3.5 | 14.4 ± 1.18 | 14.4 ± 1.18 | 2 |
| H3 1x601 BB MN | 6.87 ± 2 | 48.3 ± 3.9 | 66 ± 1.6 | 34 ± 1.6 | 9.6 ± 0.4 | 9.6 ± 0.4 | 2 |
| CA 1x601 BB | 6.8 ± 2 | 50.21 ± 6.8 | 51 ± 10 | 49 ± 10 | 6.6 ± 1.6 | 6.6 ± 1.6 | 2 |
| **Chromatin** | | | | | | | |
| H3 DA3 (1xBB) | 3.6 ± 2.0 | 17.6 ± 6.7 | 47 ± 22 | 52 ± 22 | 5.7 ± 1.3 | 0.48 ± 0.11 | 3 |
| CA DA3 (1xBB) | 4.9 ± 0.46 | 51.39 ± 5.3 | 67 ± 2.7 | 33 ± 2.7 | 14 ± 3.2 | 1.2 ± 0.27 | 4 |
| **DNA -CENP-B 1-150** | | | | | | | |
| 1x 601 | 0.9 ± 0.06 | 5.2 ± 0.9 | 59 ± 4 | 41 ± 4 | 9.2 ± 1.8 | 9.2 ± 1.8 | 4 |
| 1x601 BB | 2.6 ± 0.15 | 22.82 ± 4.4 | 71 ± 0.03 | 29 ± 0.03 | 11.1 ± 3.1 | 11.1 ± 3.1 | 4 |

**Table S5 | All kinetic parameters of CENP-B interacting with DNA, nucleosomes and chromatin.** Under the measurement conditions, ?? was determined to be 168 s[6]. Reported rate constants are not further corrected for photobleaching.

### Table S6

| | | CENP-A DA1BB FRET | | | CENP-A DA3BB FRET | | H3 DA1BB FRET | | H3 DA3BB FRET | |
|---|---|---|---|---|---|---|---|---|---|---|
| **CENP-B conc.** | | **500 pM** | **5 nM** | **50 nM** | **5 nM** | **50 nM** | **5 nM** | **50 nM** | **5 nM** | **50 nM** |
| **LF** | **A1** | 5.5 ± 0.26 | 5.98 ± 0.79 | 5.07 ± 2.16 | 5.5 ± 0.05 | 5.11 ± 0.56 | 2.74 ± 0.53 | 3.92 ± 0.12 | 1.72 ± 0.03 | 1.54 ± 0.89 |
| | **c1** | 0.02 ± 0.02 | 0.008 ± 0.002 | 0.06 ± 0.05 | 0.08 ± 0.04 | 0.06 ± 0.05 | 0.05 ± 0.04 | 0.05 ± 0.002 | 0.11 ± 0.01 | 0.08 ± 0.01 |
| | **s1** | 0.08 ± 0.004 | 0.08 ± 0.014 | 0.12 ± 0.05 | 0.08 ± 0.01 | 0.074 ± 0.008 | 0.14 ± 0.04 | 0.12 ± 0.001 | 0.118 ± 0.002 | 0.1 ± 0.02 |
| | **% area** | 56.9 ± 6.3 | 65.87 ± 2.02 | 76.79 ± 4.7 | 57.9 ± 8.61 | 49.17 ± 0.19 | 46.07 ± 3.4 | 59.64 ± 2.12 | 25.64 ± 1.49 | 19.4 ± 7.4 |
| **HF** | **A2** | 1.95 ± 0.18 | 1.65 ± 0.04 | 1.11 ± 0.35 | 2.93 ± 0.09 | 3.73 ± 0.09 | 3.24 ± 0.17 | 2.30 ± 0.4 | 6.62 ± 0.36 | 6.66 ± 0.21 |
| | **c2** | 0.35 ± 0.01 | 0.42 ± 0.04 | 0.54 ± 0.2 | 0.31 ± 0.03 | 0.24 ± 0.06 | 0.54 ± 0.05 | 0.53 ± 0.015 | 0.46 ± 0.04 | 0.096 ± 0.003 |
| | **s2** | 0.17 ± 0.044 | 0.16 ± 0.003 | 0.16 ± 0.01 | 0.11 ± 0.027 | 0.10 ± 0.002 | 0.13 ± 0.013 | 0.14 ± 0.03 | 0.09 ± 0.008 | 0.443 ± 0.008 |
| | **% area** | 43.1 ± 6.3 | 34.12 ± 2.02 | 23.2 ± 4.7 | 42.09 ± 8.61 | 50.82 ± 0.19 | 53.92 ± 2.02 | 40.35 ± 2.12 | 74.35 ± 1.49 | 80.55 ± 7.4 |
| **N total traces** | | 219 | 191 | 196 | 354 | 304 | 664 | 388 | 289 | 248 |
| **N of repeats** | | 2 | 2 | 2 | 2 | 2 | 3 | 2 | 2 | 2 |

**Table S6 | Summary of Gaussian fits from chromatin remodeling induced by CENP-B invasion experiments.**

### Table S7

| Description | Sequence |
|---|---|
| hs.Ri.CENPA.13.1-SEQ1 | rGrUrUrCrUrGrGrUrUrArCrUrUrCrUrArGrUrArArArUrUCC |
| hs.Ri.CENPA.13.1-SEQ2 | rGrGrArArUrUrUrArCrUrArGrArArGrUrArArCrCrArGrArArCrArU |
| hs.Ri.CENPA.13.2-SEQ1 | rGrArUrGrUrUrCrUrGrGrUrUrArCrUrUrCrUrArGrUrArAAT |
| hs.Ri.CENPA.13.2-SEQ2 | rArUrUrUrArCrUrArGrArArGrUrArArCrCrArGrArArCrArUrCrArA |
| hs.Ri.CENPA.13.3-SEQ1 | rArCrArArGrGrUrUrGrGrCrUrArArArGrGrArGrArUrCrCGA |
| hs.Ri.CENPA.13.3-SEQ2 | rUrCrGrGrArUrCrUrCrCrUrUrUrArGrCrCrArArCrCrUrUrGrUrCrU |

**Table S7 | Sequences of all siRNA oligonucleotides**

### Supplementary Figures

#### Figure S1

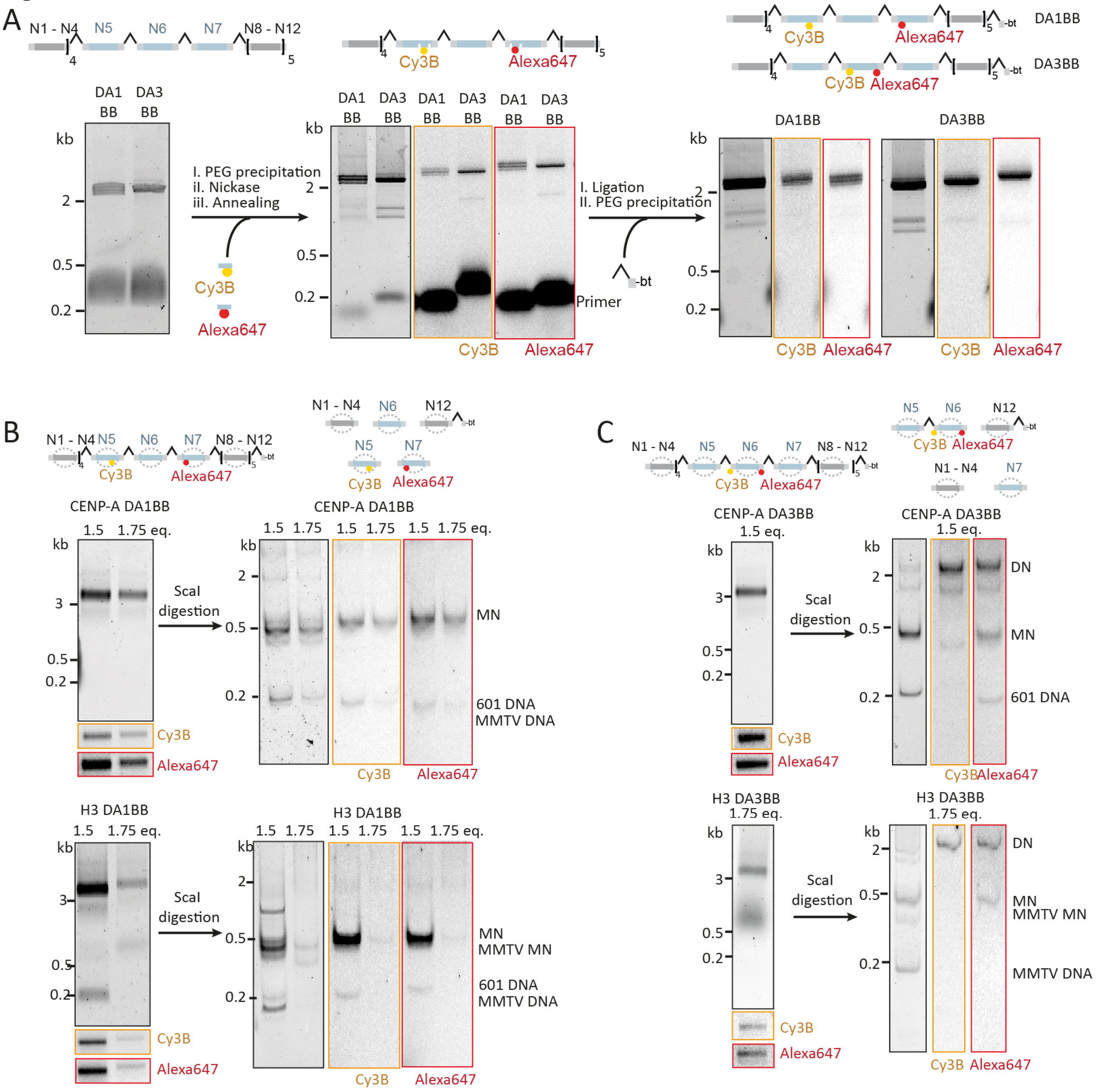


**Figure S1 - Chromatin assembly. A**) Scheme and native PAGE analysis of DNA assembly containing FRET donor (Cy3B, yellow) and acceptor (Alexa fluor 647, red) at nucleosome N5 and N7 (DA1) and at N6 (DA3) on B-Box containing (BB) DNA using plug-and-play DNA ligations[9]. 12 x 601 NPS containing DNA is purified and nicked by nicking endonucleases to create gaps into which fluorophore containing primers are annealed. All gels are imaged using GelRed, and fluorescence imaging for Cy3B and Alexa647. For DNA constructs, see **Tables S1-S3. B**) Preparation of CENP-A (top) and H3 (bottom) containing chromatin arrays on DA1BB DNA template. Saturated arrays are shown at 1:1.75 DNA to histone octamer ratio for CENP-A and 1:1.5 DNA to octamer ration for H3. Saturation is confirmed by ScaI digestion and SDS PAGE analysis of the digestion products. MMTV DNA is used for H3 as a low affinity histone buffer. For DNA constructs, see **Tables S1-S3. C**) Preparation of CENP-A (top) and H3 (bottom) containing DA3BB chromatin arrays. Saturated arrays are shown at 1:1.5 DNA to octamer ratio for CENP-A and 1:1.75 DNA to octamer ration for H3. Saturation is confirmed by ScaI digestion. As the ScaI site is obstructed by the cy3B fluorophore we see most labelled nucleosomes as dinucleosomes. For DNA constructs, see **Tables S1-S3**

### Figure S2

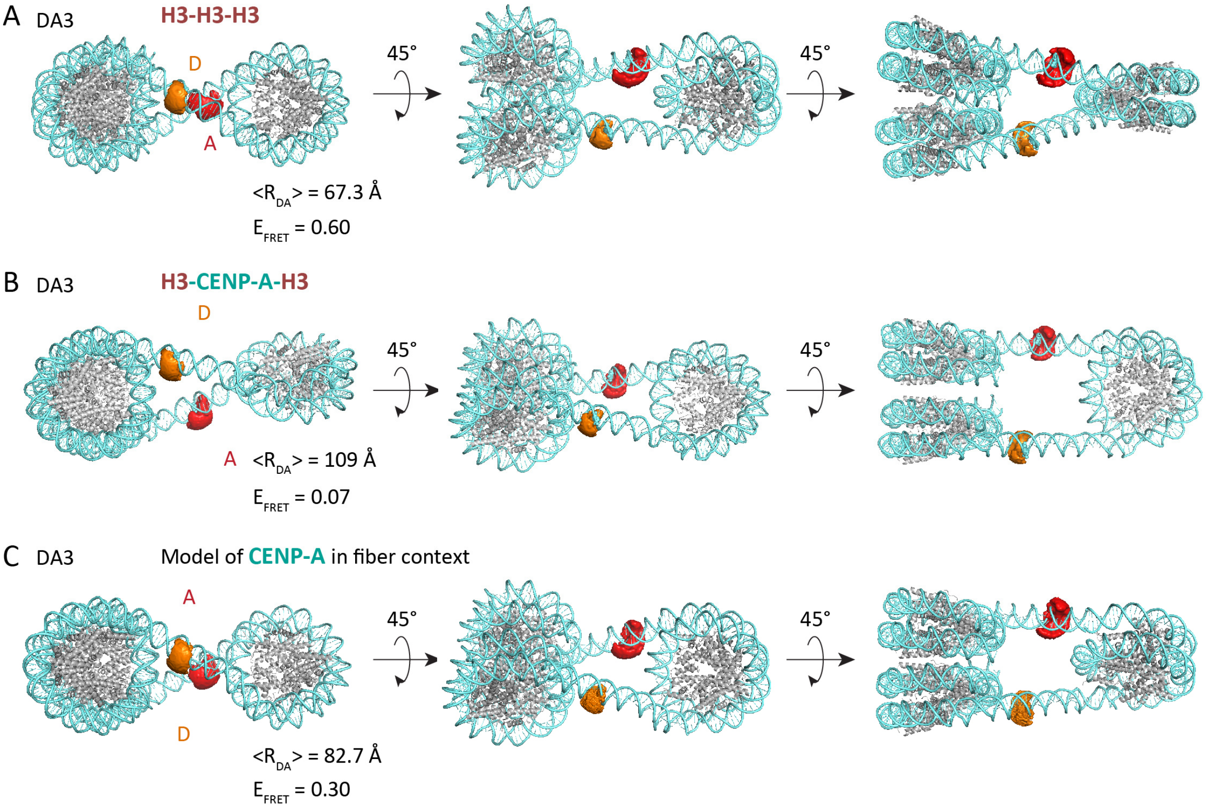


**Fig S2 – Calculated inter-dye distances and $E_{FRET}$ values for DA3 H3 and CENP-A chromatin structures. A**) H3-H3-H3 trinucleosome structure from ref. [10], PDB code 6l4a, with DA3 FRET pair, in the DNA 'outward path' orientation. Inter-dye distances based on dye accessible volumes (AV) were calculated using the FRET positioning and screening toolkit[11]. Calculated $E_{FRET}$ values ($E_{FRET}$ = 0.6) are comparable to experimentally observed values for H3 ($E_{FRET}$ = 0.48) chromatin fibers (see **Figure 2**). **B**) H3-CENP-A-H3 trinucleosome structure from ref. [10], PDB code 6l49. The calculated inter-dye distance $\langle R_{DA} \rangle$ = 109 Å is too large to obtain measurable FRET in this conformation. **C**) Chromatin state compatible with observed FRET values. Here, the conformation lies in between a H3-H3-H3 and H3-CENP-A-H3 array, with an intermediate twist of the central nucleosome, resulting in a calculated $E_{FRET}$ value of 0.3, close to the observed value of $E_{FRET}$ values = 0.32.

**Figure S3**

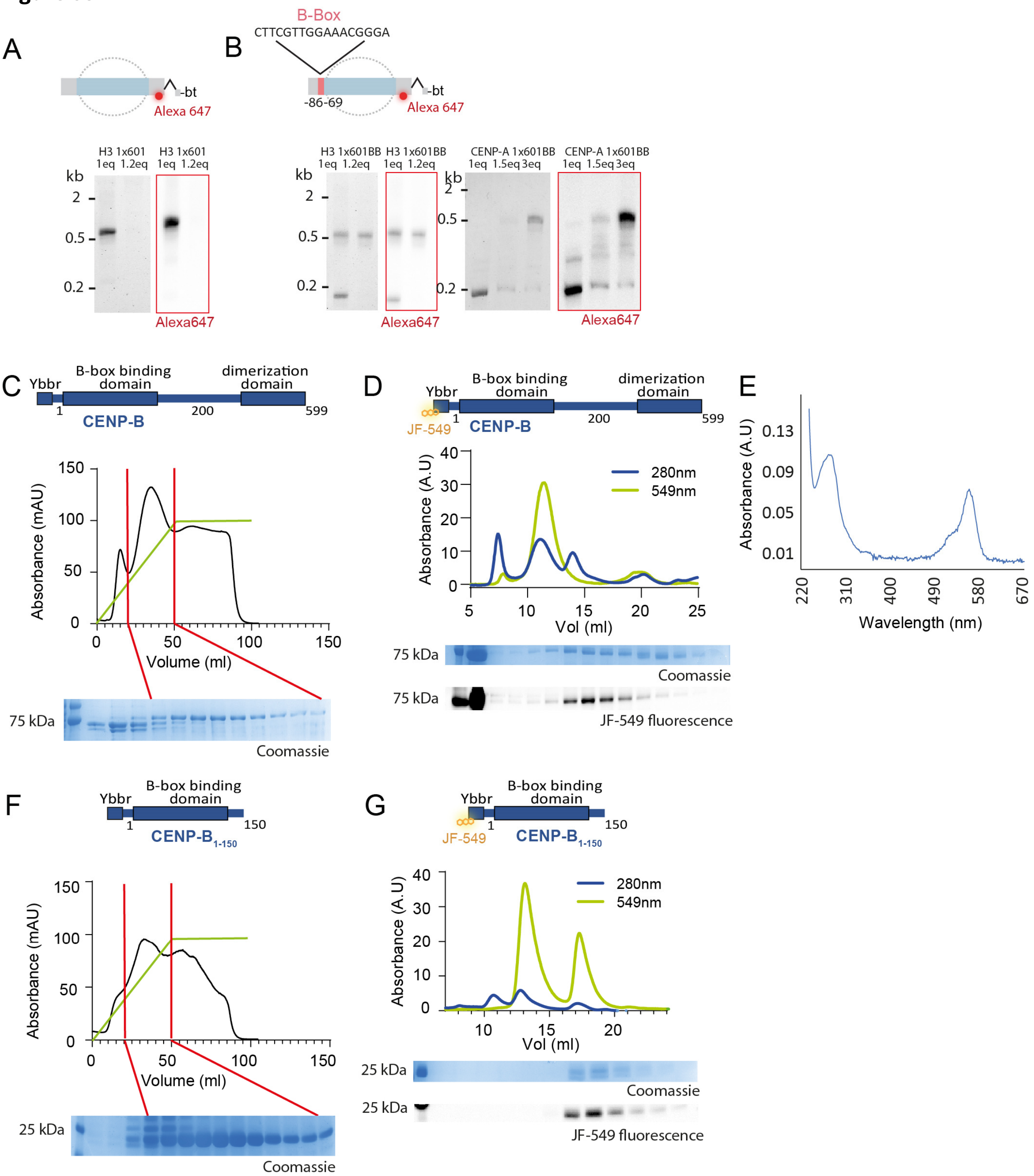


**Figure S3 - Mononucleosome formation and CENP-B production. A**) Scheme and native PAGE analysis of H3 mononucleosome assembly on DNA lacking a B-Box. For the sequence and labeling see **Tables S1-3**. **B**) Scheme and native PAGE analysis of H3 and CENP-A mononucleosome assembly on DNA with a positioned B-Box. For the sequence and labeling see **Tables S1-3**. **C**) Gradient Ni-NTA affinity purification of CENP-B (left) and CENP-B 1-150 (right). **D**) Purification of labeled of CENP-B. Gel filtration profile shows excitation spectra at 280 nm and emission spectra at 549 nm. Gels show Coomassie (top) and 549nm emission (bottom) lane 1 as input and lane 2-13 as elute of peaks 1, 2, &3 each being void, CENP-B, and sfp synthase. **E**) UV-Vis spectrum of labeled CENP-B, enabling a quantification of labeling efficiency (50%). **F**) Gradient Ni-NTA affinity purification of CENP-B(1-15). **G**) Purification and labelling of JF549-CENP-B(1-150). Gel filtration profile shows excitation spectra at 280nm and emission spectra at 549nm. Gels show Coomassie (top) and 549nm emission (bottom) lane 1 as input and lane 2-13 as elute of peaks 1, 2, &3 each being sfp synthase, CENP-B 1-150, and free dye.

**Figure S4**

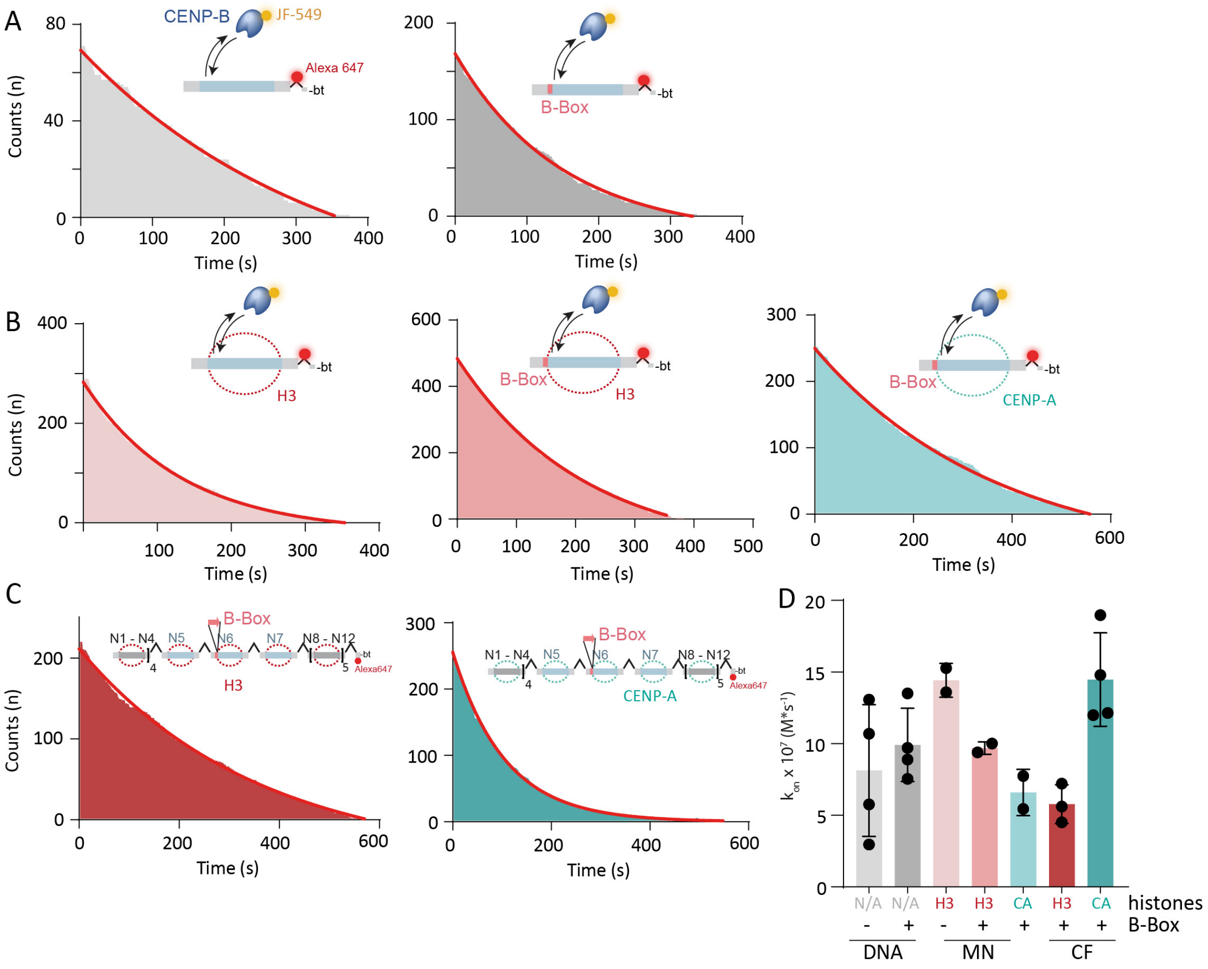


**Figure S4 - Binding kinetics of CENP-B. A**) Cumulative histogram of dark times ($t_{dark}$), yielding binding kinetics, and mono-exponential fits of CENP-B binding to 1x601 and 1x601 BB DNA. **B**) Example of cumulative histogram of dark times and mono-exponential fits of CENP-B binding to H3 1x601, 1x601 BB mononucleosomes, and CENP-A 1x601BB mononucleosomes. **C**) Example of cumulative histogram of dark times and mono-exponential fits of CENP-B binding to H3 or CENP-A containing 12x601 DA3BB chromatin arrays. **D**) Specific association kinetics of CENP-B to indicated DNA, nucleosomes (MN) or chromatin fibers (CF).

**Figure S5**

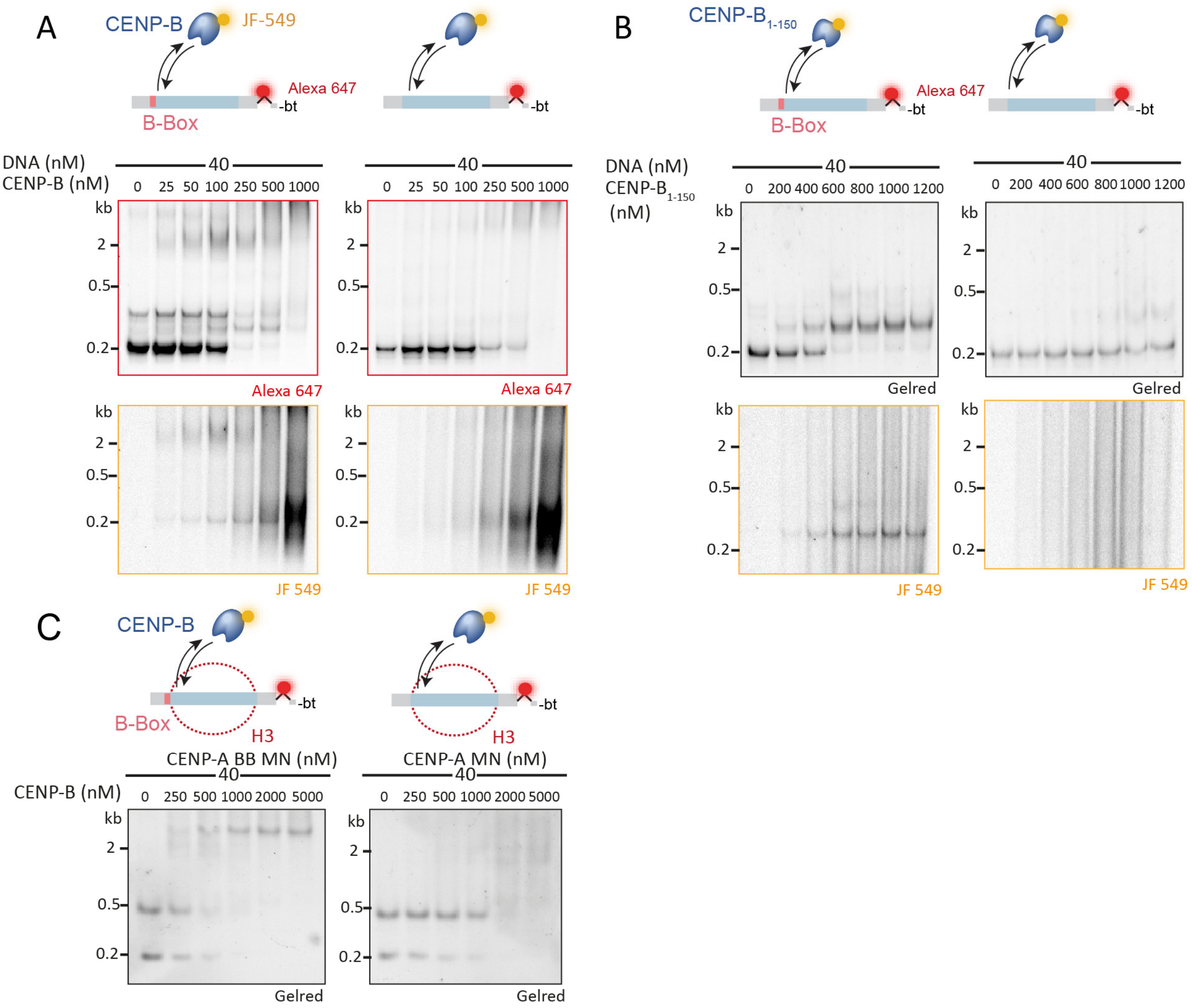


**Figure S5 – CENP-B DNA and nucleosome binding. A**) CENP-B binding to B-Box (left) and non B-Box (right) containing 1x601 DNA. CENP-B is titrated from 0-1 µM. **B**) CENP-B 1-150 binding to B-Box (left) and non B-Box (right) containing 1x601 DNA. CENP-B is titrated from 0-1.2 µM. **C**) CENP-B binding to B-Box (left) and non B-Box (right) containing 1x601 CENP-A mononucleosomes. CENP-B is titrated from 0-5 µM.

**Figure S6**

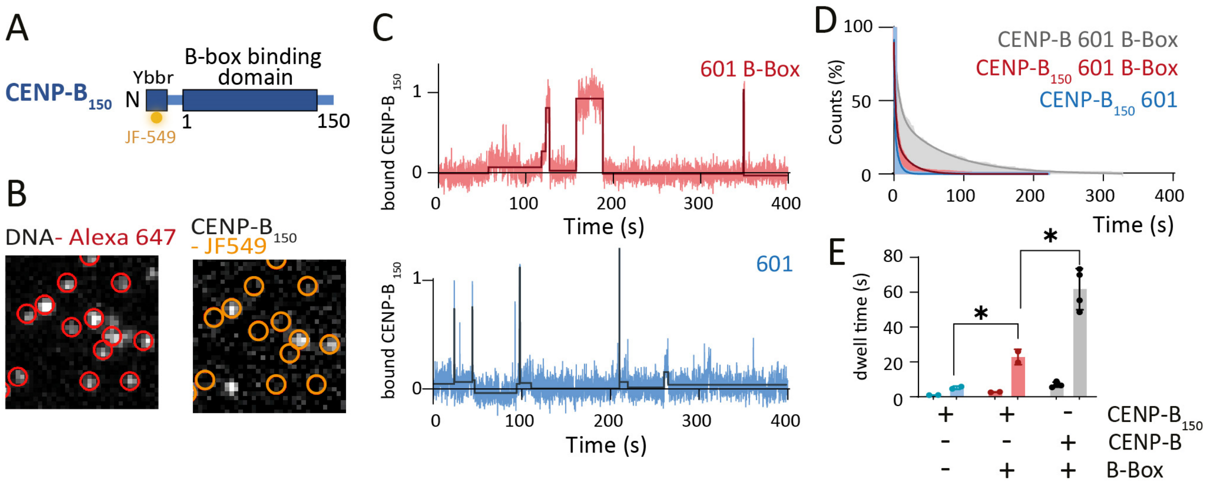


**Figure S6: CENP-B kinetochore binding and dimerization domain enhance binding to B-Box DNA. A**) Domain organization of CENP-$B_{150}$ **B**) Representative smTIRF image showing immobilized 601-B-Box DNA in the far-red channel (left, red circles) and CENP-B binding events in the green-orange channel at the same positions (right, orange circles). **C**) Representative fluorescence time trace of CENP-B 1-150 binding events to 601-B-Box (top) and non-B-Box DNA (bottom). The traces are fitted and $t_{dark}$ and $t_{bright}$ are determined. **D**) Cumulative histogram of CENP-B 1-150 binding to B-Box (red) and non-B-Box DNA (blue), and CENP-B Fl binding to 601 B-Box DNA (grey) fitted by a bi-exponential function (solid line). **E**) Specific disassociation time constants ($t_{off,i}$) of CENP-B 1-150 to non-B-Box (blue) and B-Box (red) 601 DNA, and CENP-B Fl to B-Box 601 DNA. n=2-4; error bars are S.D. *<0.05 using 2-tailed unpaired t-test.

**Figure S7**

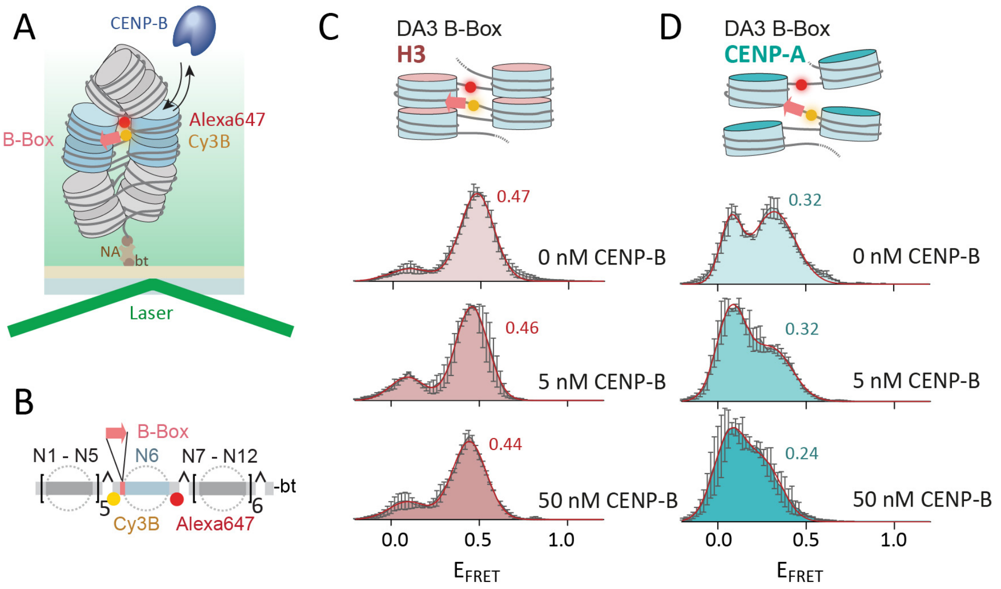


**Figure S7 – Effect of CENP-B binding on linker DNA orientation within chromatin. A**) Scheme of smFRET – TIRF experiment using a DA3BB DNA template. bt is biotin, NA is neutravidin. **B**) Scheme of the used chromatin DNA containing FRET donor (Cy3B, yellow) and acceptor (Alexa fluor 647, red) and 1xB-Box at N6 (DA3BB). **C**) DA3 FRET histograms for H3 DA3BB at indicated CENP-B concentrations. Histograms are fitted with Gaussian functions (red), and the $E_{FRET}$ value of the high FRET population is displayed. **D**) CENP-A DA3 histograms at indicated CENP-B concentrations with the $E_{FRET}$ value of the high FRET population displayed. All Histograms are the average of n=2 independent repeats. Peak $E_{FRET}$ for high FRET populations are indicated. Error bars are standard error.

**Figure S8**

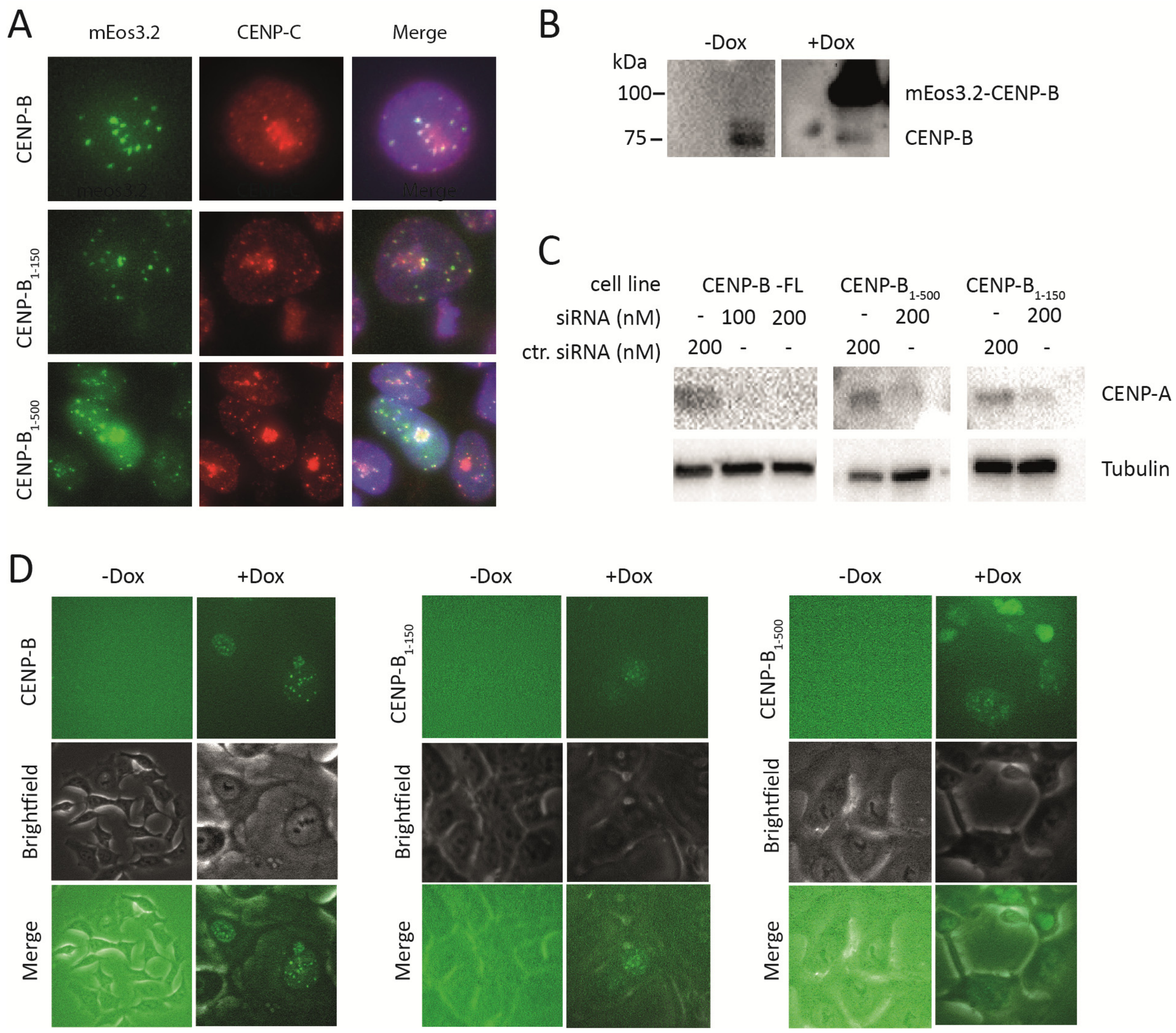


**Figure S8 - CENP-B mutant cell line preparation. A**) Immunofluorescence images showing colocalization of meos3.2-CENP-B and CENP-C puncta at centromeres. **B**) Western blot against CENP-B for meos3.2-CENP-B expressing cells before and after doxycycline addition. CENP-B1-150 and 1-500 do not have the required epitope for our CENP-B antibody. **C**) Western blot analysis of siRNA knockdown of meos3.2-CENP-B fl/1-500/1-150. **D**) Live cell imaging of meos3.2-CENP-B fl/1-500/1-150 before and after doxycycline addition.

**Figure S9**

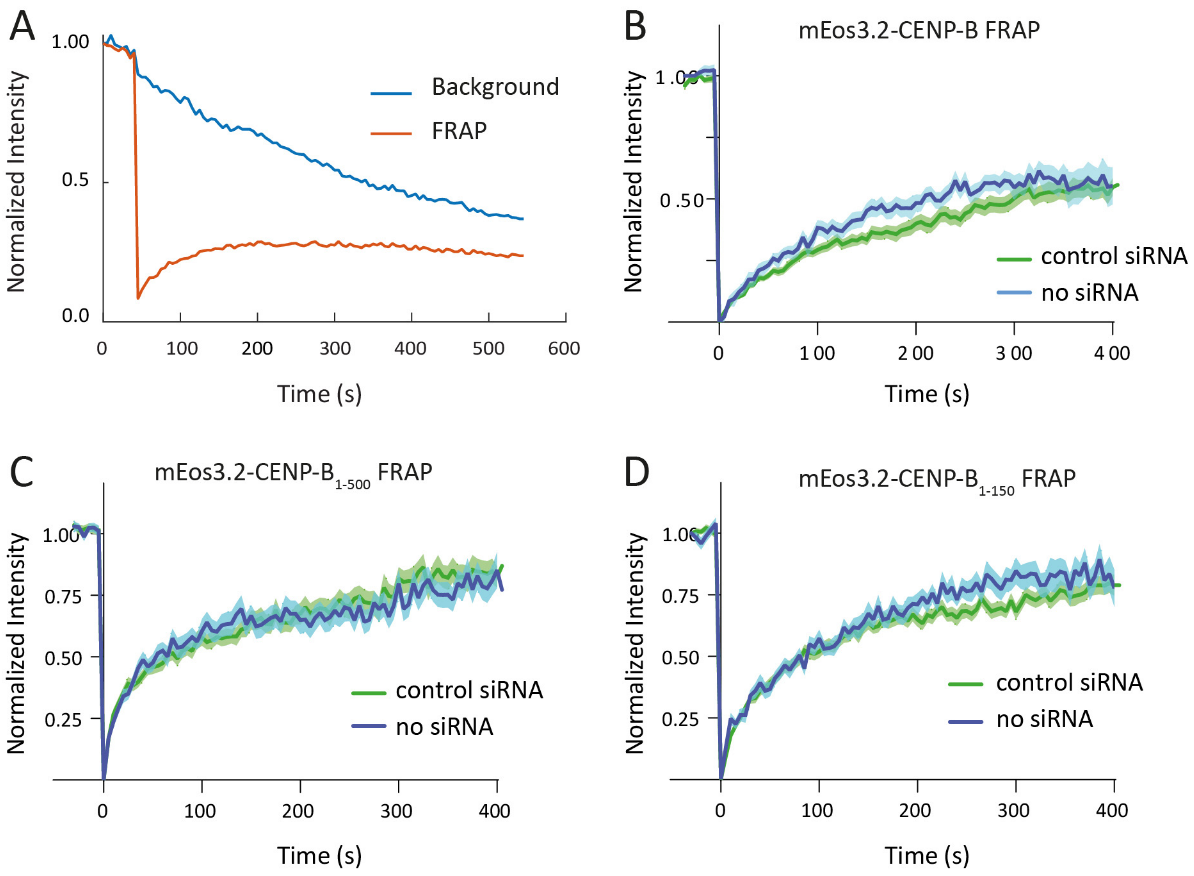


**Figure S9 – FRAP analysis of CENP-B binding. A**) Example raw FRAP curve for meos3.2-CENP-B comparing intensity of unbleached vs bleached CENP-B puncta. **B**) FRAP curve for meos3.2-CENP-B-Fl without siRNA treatment (blue) and negative control siRNA transfection (green). Error bars in SEM. **C**) FRAP curve for mEos3.2-CENP-B-1-500 without siRNA treatment (blue) and negative control siRNA transfection (green). Error bars in SEM. **D**) FRAP curve for meos3.2-CENP-B-1-150 without siRNA treatment (blue) and negative control siRNA transfection (green). Error bars in SEM.

**Figure S10**

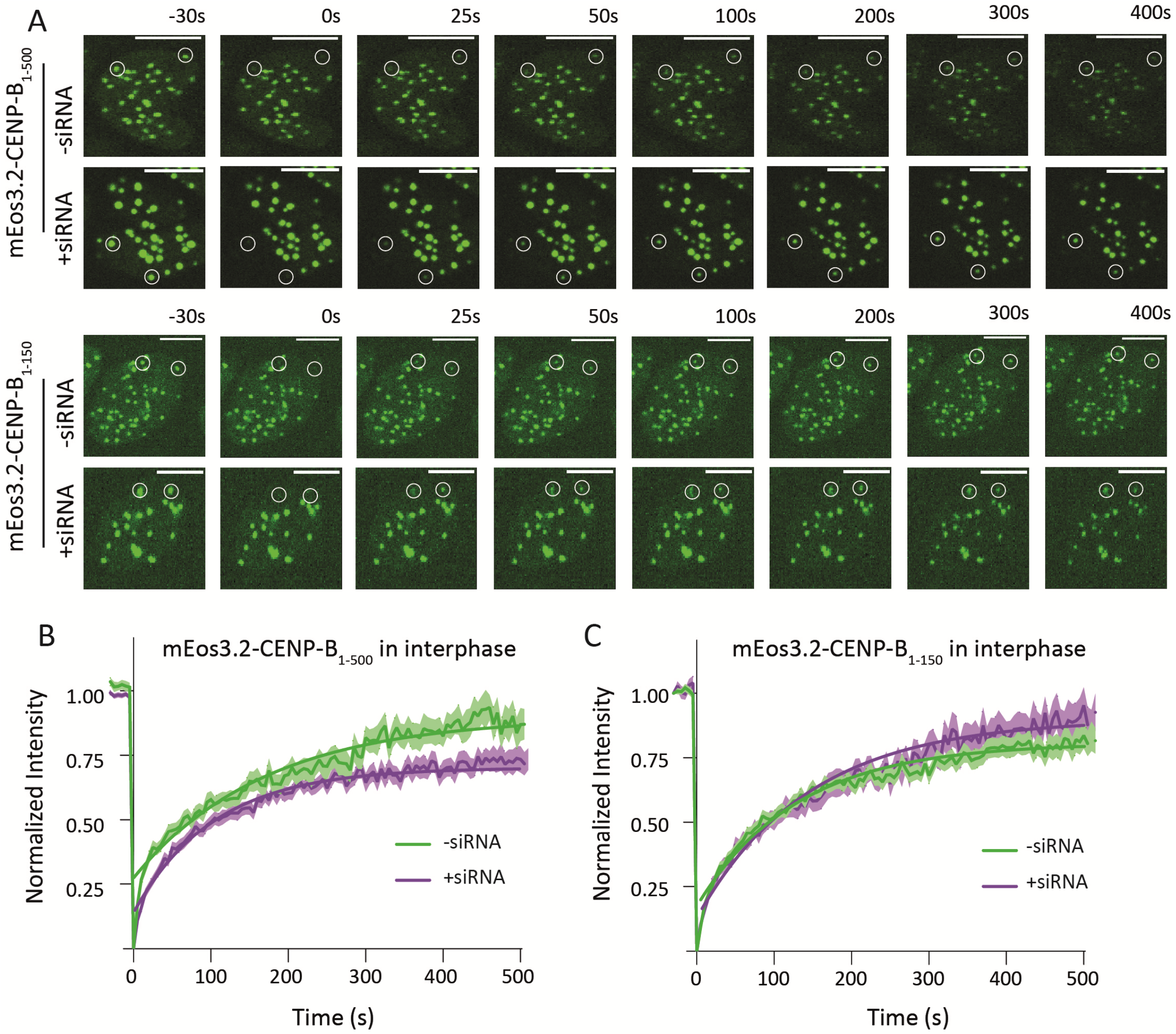


**Figure S10: FRAP analysis of indicated CENP-B mutants with and without CENP-A depletion in interphase cells. A**) Representative FRAP analysis of mEos3.2-CENP-B 1-500 (top) an mEos3.2-CENP-B 1-150 (bottom) in the presence of CENP-A (top, -siRNA) or absence of CENP-A (bottom, 48hrs after siRNA treatment). Bleached area is indicated with white circles. T<0 frame is before bleaching and signal recovery is shown at indicated time points. **B**) Normalized recovery curves for quantitative FRAP measurements of CENP-B 1-500 with or without CENP-A depletion. Thick lines indicate mean values and the colored area between the thin lines indicated the standard error. At least 10 cells and 20 centromeres were used for analysis. Exponential fitting yielded a time constant of recovery of $\tau$ = 171 ± 11 s (- siRNA) and $\tau$ = 104 ± 3.5 s (+ siRNA). **C**) Normalized recovery curves for quantitative FRAP measurements of CENP-B 1-150 with or without CENP-A depletion. Thick lines indicate mean values and the colored area between the thin lines indicated the standard error. At least 10 cells and 20 centromeres were used for analysis. Exponential fitting yielded a time constant of recovery of $\tau$ = 128 ± 6 s (- siRNA) and $\tau$ = 144 ± 7 s (+ siRNA).
